# Evaluation of extraction solvents for untargeted metabolomics to decipher the dissolved organic matter of Antarctic cryoconite holes

**DOI:** 10.1101/2024.04.29.591772

**Authors:** Swapnil Mundhe, Saborni Maiti, Aritri Sanyal, Narendra Y Kadoo, Dhiraj Dhotre, Vitthal T Barvkar, Shamim A. Shaikh, Runa Antony, Dhiraj Paul

## Abstract

Cryoconite holes are biological hotspots with a high biogeochemical turnover rate, contributing significantly to the glacial ecosystem’s overall carbon cycles and net fluxes. Unfortunately, the information about the composition of low molecular weight molecules formed through the metabolic processes of cryoconite-dwelling microbes is scanty. These molecules constitute a substantial portion of the dissolved organic matter (DOM) within cryoconite holes. The present study investigated the composition of DOM in cryoconite holes using reverse-phase liquid chromatography (RP-LC) coupled with high-resolution tandem mass spectrometry. We evaluated various solvent combinations of water, methanol, and acetonitrile to extract chemically diverse polar and non-polar metabolites from the cryoconite holes. Among the single solvents, organic-rich MeOH: Water (70:30 v/v) and in parallel 2-single solvent combinations of MeOH: Water (70:30 v/v) and Acetonitrile: Methanol: Water (40:40:20 v/v) provided increased number and chemical diversity of extracted metabolites. Combining RP with the hydrophilic interaction liquid chromatography (HILIC) technique provided the highest number of unique metabolites. This dual-LC and ionization polarity combination increased the detection of metabolic features by 46.96% and 24.52% in single- and two-solvent combinations compared to RP alone. This study developed a simple untargeted metabolomics workflow that is highly sensitive and robust, detecting and potentially identifying a large number of chemically diverse molecules present in the DOM (extracellular) and microbes (intracellular) from the cryoconite holes environment. This method can better characterize DOM’s chemical composition and, after integrating with other ‘omics’ approaches, can be used to examine the link between metabolic pathways and microbial communities in global cryoconite holes or other similar ecosystems, revealing how these earthy systems and their microbial flora control carbon or nutrient storage or release in response to global climate change. Overall, the study presents a valuable methodology for studying the biogeochemistry of cryoconite holes.

## 1. Introduction

Microorganisms are a critical component in the terrestrial biosphere as they are actively involved in the cycling of the organic compounds in dissolved organic matter (DOM) by synthesis and decomposition of microbial metabolic products (Schmidt et al., 2022). Low molecular weight (LMW, 50-1500 Da) molecules, such as small peptides, carbohydrates, nucleic acids, organic acids, lipids, and other metabolites in the DOM fractions, are readily available for microbial processing (Ladd et al., 2019). Therefore, fingerprinting of LMW molecules in the DOM provides a detailed snapshot of organic molecules consumed or produced by microbes. This is especially relevant for Antarctic ecosystems that are experiencing rapid changes due to climate change. However, the heterogeneity of this pool, coupled with consistently low concentrations, poses significant challenges in detecting and quantifying these molecules in the Antarctic or other such extreme ecosystems.

Cryoconite holes (CHs) are self-contained water-filled cylindrical holes with cryoconite (dark-colored sediment) deposited at their undermost and are formed on ice sheets and glaciers (Poniecka et al., 2020; Rozwalak et al., 2022) covering 1–20% of the world’s ice (Hodson et al. 2008). They impact the ablation process of glacial ice and serve as the habitats of glacial microbes (Bagshaw et al. 2013; Zawierucha et al. 2018). They are formed when organic or inorganic particles are deposited on the glacial ice surface. Being darker, they absorb more solar energy than the surrounding ice, melting it and creating water-filled cylindrical depressions (Gribbon, 1979; Cook et al., 2016). Because of dark mineral and organic/inorganic matter (cryoconite) at the bottom covered by ice melt-water, CHs can have a much higher biogeochemical turnover rate than in other supraglacial zones (Bagshaw et al., 2013; Zawierucha et al., 2018a, 2018b, and 2022). Despite their small individual size but very high numbers, the occurrence of several microbial-mediated biogeochemical processes within them can significantly contribute to the glacier ecosystem’s overall nutrient cycles and net fluxes (Anesio et al., 2009 and 2017). Due to this unique nature of the high content of DOM and organic compounds, CHs allow diverse microbial communities to flourish in this unique and harsh environment (Sanyal et al., 2018; Sommers et al., 2019). While several studies have reported microbial diversity and its composition in the CHs ecosystem, only a few have applied metabolomics to reveal the metabolic potential of the indigenous microbial communities of this ecosystem in the Greenland ice sheet and Arctic ice cap (Cook et al., 2016; Gokul et al., 2023; Doting et al., 2024). Similarly, Lutz et al. (2015) employed metabolomics to assess the differences between a red and a green snowfield on a glacier in Svalbard, Norway. Despite the importance of microbial metabolites in nutrient cycles and ecosystem dynamics, optimizing an untargeted metabolomics approach to investigate the LMW DOM and their roles in the CHs systems of either the northern or southern hemispheres is very limited.

Untargeted metabolomics is a robust and rapidly growing approach to analyzing complex extracts by providing comprehensive data-driven metabolism analyses (Di Minno et al., 2021; Bittremieux et al., 2023; Bowen et al., 2023). Several studies have employed the untargeted metabolomics approach to reveal a wide range of compounds in the DOM of terrestrial and marine ecosystems. This has improved our understanding of the significance of DOM in structuring microbial populations and its role in global climate change. These studies are gaining much recent interest and giving rise to an emerging area of study (Swenson et al., 2015; Ladd et al., 2019, 2021; Catalá et al., 2021; Yao et al., 2023). Glacial ecosystems like CHs can produce significant amounts of DOM because of the biochemical activities, which can potentially seed this bio-available material to nearby aquatic environments (Foreman et al., 2007; Feng et al., 2016). Although the high DOM and other nutrients are known to be present in CHs, the interactions between various organic substrates and microbes in the CHs are poorly understood (Weisleitner et al., 2020; Sanyal et al., 2024).

While some studies have explored DOM characterization at the compound level in various terrestrial ecosystems (Mann et al., 2012; Singer et al., 2012; Nebbioso and Piccolo et al., 2013; Pautler et al., 2013; Antony et al., 2014; Mann et al., 2015; Swenson et al., 2015; Grewer et al., 2016), the number of such investigations remains limited. In these studies, Nuclear Magnetic Resonance (NMR) spectroscopy, Fourier Transform Ion Cyclotron Resonance Mass Spectrometry (FTICR-MS), UV-visible or Excitation-Emission Matrix Fluorescence Spectroscopy and gas chromatography-mass spectrometry (GC-MS) measurements were used to apply the untargeted metabolomics approach to reveal a range of DOM compounds. Even though the choice of analytical platform is study-specific, these techniques have inherent limitations. For example, FTICR-MS can offer the highest resolution and mass accuracy but at the expense of longer runtimes due to low data acquisition rates. Thus, this approach is less suitable for high-throughput data acquisition than the orbitrap instruments (Hohenester et al., 2020). Likewise, the UV-visible or excitation-matrix spectroscope requires detectable chromophores in each analyte, and highly complex mixtures such as metabolic extracts may contain molecules that lack chromophores (Vitale et al., 2024). Similarly, NMR is less sensitive, and the number of metabolites detected is also low. In contrast, GC-MS-based experiments require volatile analytes or prior derivatization of samples (Martin et al., 2014; Ladd et al., 2019).

Consequently, in recent years, LC-ESI-MS has emerged as a powerful tool for analyzing mixtures of small molecules. Its broad detection range, high-throughput capabilities, and sensitivity make it an ideal alternative to the traditional methods of characterizing DOM (Patti, 2011; Dunn et al., 2013; Chetwynd and David, 2018). Although reverse-phase-liquid chromatography (RP-LC) in positive ionization mode is preferred, drawbacks of using only a single chromatography phase or polarity mode are reported (Gika et al., 2014). The hydrophilic interaction liquid chromatography (HILIC) technique is more suitable for retaining and separating highly polar molecules than RP-LC, which can efficiently retain and separate mid- and non-polar molecules (Boudah et al., 2014; Contrepois et al., 2015). Thus, combining RP-LC and HILIC-LC-MS techniques is an attractive complementary approach to identifying a broad spectrum of the metabolome (Tang et al., 2016). This untargeted global metabolite profiling would bridge the knowledge gap that limits our understanding of DOM composition and metabolic pathways in microbial communities (Swenson et al., 2015; Ladd et al., 2019).

In this study, we performed a comparative evaluation of the different single, binary, and ternary combinations of the most commonly used MS-grade solvents for RPLC-MS/MS-based untargeted metabolomics (positive and negative ESI mode) to extract and detect a broad range of metabolites (intra- and extracellular) from CHs samples collected from Larsemann Hills, Antarctica. Further, we combined the results with HILIC-MS/MS in ESI (+ve) and ESI (-ve) ionization modes to increase the metabolite detection range. To our knowledge, this is the first time a systematic optimization study has delineated an LC- MS/MS-based untargeted metabolomics approach to uncover DOM composition in cryoconite holes. This study lays the technical foundation for future experiments aiming to perform untargeted metabolomics analyses to understand the LMW metabolites composition of the cryoconite holes worldwide and similar ecosystems, aiming to integrate LMW DOM molecular data into process-based ecological models.

## 2. Materials and Methods

### 2.1 Chemicals

The LC-MS grade solvents methanol (CAS 67-56-1), acetonitrile (CAS 75-05-8), and water (CAS 7732-18-5) were procured from JT Baker (New Jersey, USA). Similarly, the LC-MS grade chemicals ammonium formate (CAS 540-69-2) and formic acid (CAS 64-18-6) were produced from Sigma-Aldrich (St. Louis, USA). The authentic metabolic standards such as L-alanine, L-glutamic acid, L-methionine, L-tryptophan, L-tyrosine, L-phenylalanine, L- proline, L-malic acid, L-isoleucine, L-leucine, L-valine, L-histidine, L-asparagine, L- cysteine, L-threonine, L-glutamine, L-serine, L-glutathione, D-mannitol, D-sucrose, pyridoxine hydrochloride, riboflavin were also procured from Sigma-Aldrich (St. Louis, USA).

### 2.2 Cryoconite hole sample collection and processing

Sediment samples from cryoconite holes in the Larsemann Hills region of East Antarctica were obtained during the 38^th^ Indian Scientific Expedition to Antarctica during the austral summer of 2019-20 (69°25’13.09” S, 76° 13’ 22.85” E). Dissolved oxygen, pH, temperature, oxidation-reduction potential, salinity, total dissolved solids, and conductivity were measured during sample collection (Table S1). The frozen samples were initially transported to the National Centre for Polar and Ocean Research (Vasco-Da-Gama, India) for sub-sampling and subsequently transferred to the National Centre for Cell Science, Pune, India, under dry ice and stored at -80°C until analysis. The samples were lyophilized before metabolite extraction, and coarse sand and fine gravel were removed. The samples were pulverized with a sterile, liquid N_2_ chilled mortar and pestle, and the powdered samples were stored at -80°C till metabolites isolation.

### 2.3 Metabolites extraction conditions for the cryoconite hole sediment

An equal amount of homogenized powdered samples from 20 CHs were mixed and used to evaluate extraction solvents. The following extraction method was used to optimize experimental conditions of metabolite extraction for RP-LC-MS/MS analysis. Before extraction, all the LC-MS grade solvents were filtered via a 0.22 µm syringe filter (Chromatopak, India). Eight extraction solvents of single, binary, and ternary combinations (v/v) of water, methanol, and acetonitrile were prepared as 100% water (water), 30:70 methanol: water (30MW), 30:70 acetonitrile: water (30AW), 100% methanol (MeOH), 100% acetonitrile (ACN), 70:30 acetonitrile: water (70AW) and 70:30 methanol: water (70MW) and 40:40:20 acetonitrile: methanol: water (AMW). The samples were extracted in quadruplets. The workflow of sample pre-preparation, data acquisition, and analysis is shown in Workflow 1. All extractions were performed using 2 g of sample biomass (Swenson et al., 2015). A finely powdered sample was transferred into a 2 mL centrifuge tube. 1.2 mL of ice-chilled extraction solvent was added to it, followed by vortexing for 1 min to mix the sample and solvent, with subsequent homogenization in TissueLyser II (Qiagen, Hilden, Germany) and incubation (at 4°C) for 5 min, respectively. The samples were sonicated for 10 min on ice to keep the temperature low and incubated again at 4°C for 5 min, followed by vortexing for 5 min and centrifugation at 13500 rpm for 15 min at 4°C. The supernatant was collected and filtered through a 0.22 µm syringe filter. The samples were evaporated to near dryness and reconstituted to 500 µL using the respective extraction solvent. For HILIC phase LC-MS/MS analysis, a 2 g sample was extracted in water using the above-mentioned extraction method. Dried extracts were reconstituted in 500 µL of 60% (60HILIC), 75% (75HILIC), 90% (90HILIC), and 95% (95HILIC) of the acetonitrile-water solvent combination. The samples were stored at -80 °C till further use.

### 2.4 Evaluation of extraction yield

Absolute metabolites extraction yield was estimated for RP extracts by measuring the remaining dried residue after evaporating 500 μL of solvent supernatant (extract) to dryness using the Savant DNA-120 SpeedVac concentrator (Thermo Fisher Scientific, Waltham, USA). The extraction yield was determined as a percent of the initial dry weight of the sample utilized to produce 500 μL of extract. All the extracts were digitally photographed and visually inspected for color, clarity, and resemblance.

### 2.5 LC-MS/MS data acquisition

Thermo Q-Exactive-Orbitrap™ high-resolution mass spectrometer (HRMS) coupled to an Accela™ 1200 ultra-high-performance liquid chromatography (UHPLC) system (Thermo Fisher Scientific, Waltham, USA) was used for data acquisition in ESI (+ve) and ESI (-ve) modes. The heated electrospray ionization (HESI) source settings were as follows: capillary and gas temperature was 320 °C; auxiliary, sheath, and spare gas were at 45, 12, and 2 arbitrary units, respectively, in both the modes; spray capillary voltage was 4.2 kV and 3.8 kV in positive and negative modes, respectively. The tube lens was set to 45 V, and the mass scan range was set from 50-1000 m/z. The resolution power of the Orbitrap was set at 70,000. The instrument was calibrated before running samples as per the vendor’s instruction. A Hypersil GOLD C18 column (1.9 µm, 2.1mm × 150) (Thermo Fisher Scientific, Waltham, USA) was utilized for reverse phase chromatography (RP). The column oven temperature was maintained at 40 °C, and the sample manager temperature was set at 4 °C. Mobile phases A (0.1% formic acid) and B (ACN, 0.1% formic acid) were employed to separate the metabolites. Linear gradient elution was set to 20 min with a 600 µL/min flow rate. Eluent A was reduced from 98% to 60% between the start and 5 min, 40% at 7 min, 20% at 10 min, and 2% at 13 min. It was restored to its beginning condition after 16 min and held for another 4 min. For HILIC phase chromatography, a Syncronis C18 column (1.7 µm, 2.1mm × 100) (Thermo Fisher Scientific, Waltham, USA) was used and maintained at 40 °C. Mobile phase A (95% ACN, 5% 5 mM ammonium formate, 0.1% formic acid) and B (5 mM ammonium formate, 0.1% formic acid) were used to separate metabolites. The gradient was: at 0-2 min, eluent A was 95%; at 14 min, eluent A was brought to 60%; maintained until 16 min, and 95% at 16.5 min and held until 20 min before injecting the following sample. The flow rate was set at 600 µL/min. The mass spectrometer parameters were the same as the reverse phase.

Thermo Xcalibur v. 3.0.63 (Thermo Fisher Scientific, Waltham, USA) was used to record the chromatograms and spectra in profile mode. The instrument was periodically checked for ambient operating conditions. Samples were injected randomly with an injection volume of 3 µL. Pooled quality control (QC) samples were prepared by mixing all the samples to ensure quality throughout the run and the quality of the metabolic profile. Before beginning the actual sample data acquisition, a series of QC samples were injected to equilibrate and condition the system, followed by an interval of nine samples. The samples were injected randomly with an injection volume of 3 µL. QC samples were also used for the tandem MS data acquisition. In RP, MS/MS data acquisition was conducted at 40 Normalized Collision Energy (NCE) in Parallel Reaction Monitoring (PRM) mode using an inclusion list containing the mass-to-charge ratio (m/z) of detected ions and their associated retention times (RTs) acquired from prior data analysis, and in HILIC data were acquired in data-dependent MS/MS (ddMS^2^) mode with N=10.

### 2.6 LC-MS data processing and analysis

The QC Viewer function of the RawHummus tool (https://cran.r-project.org/web/packages/RawHummus/index.html), an R package, was used to evaluate data quality, instrument stability, and reproducibility (Dong et al., 2022). Raw data processing, peak picking, and conversion of raw data from profile (.raw) to centroid mode format, i.e., mzXML of both ESI (+ve) and ESI (-ve) ionization modes were performed using ProteoWizard v 3.0.21082 (Chambers et al., 2012). Peak detection, retention time correction and alignment, peak grouping, and probabilistic putative metabolite annotation were performed using the R-based ProbMetab tool (Silva et al., 2014) equipped with upstream R packages like xcms (Smith et al., 2006), CAMERA (Kuhl et al., 2012), mzMatch (Scheltema et al., 2011), and Comb2+ algorithm. The metabolic features were putatively assigned KEGG IDs and compound names at the MS1 level by searching against the KEGG (Kyoto Encyclopedia of Genes and Genomes, http://www.kegg.com) database. The different metabolic classes were assigned using the R Package ‘omu’ (Tiffany and Bäumler et al., 2019). Data for the ESI (+)ve and ESI (-)ve ionization modes were processed separately for RP and HILIC and merged for further analysis (Tables S2, S3, S4, and S5). The R packages ggplot2, reshape, and pheatmap were used to generate scatter plots, stack bar plots, and heat maps. Venn diagrams were generated using InteractiVenn, a web-based tool (Heberle et al., 2015). For the metabolic feature reproducibility, the coefficient of variation (CV<25%) was determined from the peak areas among replicates of each solvent extract in RP and HILIC. The term metabolic feature in this article refers to the chemical moiety peak detected at unique m/z and RT.

For multivariate statistical analysis of the RP data (all solvents combined), duplicate and unknown features were first removed. Artificial artifacts, noises, and contaminations were removed manually using the blanks as a reference. The resulting data frame was uploaded to the web-based platform MetaboAnalyst 5.0 (http://www.metaboanalyst.ca). The data filtering step applied a 10% variance filter based on the interquartile range (IQR) to remove detected near-constant features across all solvents. Further, the data were normalized by sum to adjust systematic differences among samples followed by log transformation (base 10) and Pareto scaling (mean-centered and divided by the square root of the standard deviation of each variable). Fig. S1 shows the data distribution pattern before and after normalization.

After this, parametric one-way ANOVA was applied to select 499 significant features (*p*<0.05, Table S6), followed by principal component analysis (PCA). For hierarchical cluster analysis (heatmap), we selected the metabolic features with variable importance in projections, VIP score >1.5 (Fig. S2) via partial least square discriminant analysis (PLS-DA, Fig. S3) (Eriksson et al., 2006). To validate the metabolite identification rigorously, the Metabolomics Standard Initiative (MSI) (Fiehn et al., 2007) recommends different levels of identification (Sumner et al., 2007). In our study, from the metabolic features obtained by PLS-DA, eight features were confirmed (MSI standards level 1), and 19 were putatively confirmed (MSI standards level 2). Confirmation of identification of the m/z was performed using MS/MS spectral match with authentic standards (Table S7 and Fig. S4). Where authentic standards were not available, putative confirmation of features was carried out by extracting the MS2 spectra (fragment list) of the parent ion (of the desired metabolic feature) using the Qual browser of Xcalibur v. 3.0.63 (Thermo Fisher Scientific, Waltham, USA) and matching with theoretically predicted fragments of the same parent ion from Mass Frontier v. 7.0 (Thermo Fisher Scientific, Waltham, USA) software (Table S8). The workflow for data analysis for RP and HILIC is shown in Workflow S1. Data were reported at levels 1,2, and 3 according to the metabolite standard initiative guidelines as mentioned above.

### 2.7 Detection of low abundance metabolic features

A cumulative abundance plot was used to evaluate the solvent extraction and chromatographic technique in extracting and detecting the least abundant metabolic features. The total number of detected features in the individual analysis of all eight solvents was used for evaluation. For RP and HILIC techniques, the highest number of detected features (i.e., 70MW and 75HILIC) were compared in both positive and negative modes.

The proportional abundance of each feature contributing to the total abundance of all metabolic features detected was calculated using the following formula:

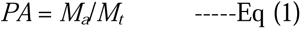

*PA*=Proportional abundance

*M_a_*=abundance of each metabolic feature

*M_t_*= total abundance of all metabolic features

Cumulative abundance was calculated using the following formula:

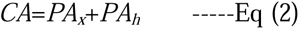

*CA*= cumulative abundance

*PA_x_*= proportional abundance of the *x^th^* rank metabolic feature

*PA_h_*=proportional abundance of metabolic feature *(x+1) ^th^* higher in rank

## 3. Results

The present study aimed to optimize and demonstrate the high-throughput, sensitive, untargeted approach to detect, quantify (relatively), and putatively annotate variations in the LMW DOM availability across space in cryoconite holes. Therefore, to enable characterization and to expand the maximum coverage of metabolites, we evaluated several extraction solvent combinations for RP-LCMS-based untargeted metabolomics of LMW compounds in CHs in combination with HILIC-LCMS data.

### 3.1 Extraction efficiencies

The extraction efficiencies of the eight solvent systems used to extract metabolites from the DOM of the CHs were evaluated (Fig. 1A). According to the percentage extraction yield, the solvent systems were grouped as high (70AW, 70MW, ACN), moderate (MeOH, AMW, Water), and low (30MW, 30AW) efficient. A visual inspection showed that all extracts were clear and transparent but varied in color (Fig. S5 A and S5 B). Aqueous and aqueous-rich solvent extracts such as water, 30AW, and 30MW were colorless, whereas organic and organic-rich solvent extracts like MeOH, ACN, 70AW, and AMW were colored (light yellow and green). Interestingly, despite 70MW being an organic-rich solvent, the extract was colorless. The percentage extraction yield ranged between 0.66 and 1.56%. The solvent 70AW showed the highest extraction yield (1.56%). Overall, across all the solvent systems, the extraction efficiency was in the order: 30AW<30 MW<Water<AMW<MeOH<ACN<70MW<70AW.

**Figure 1:**
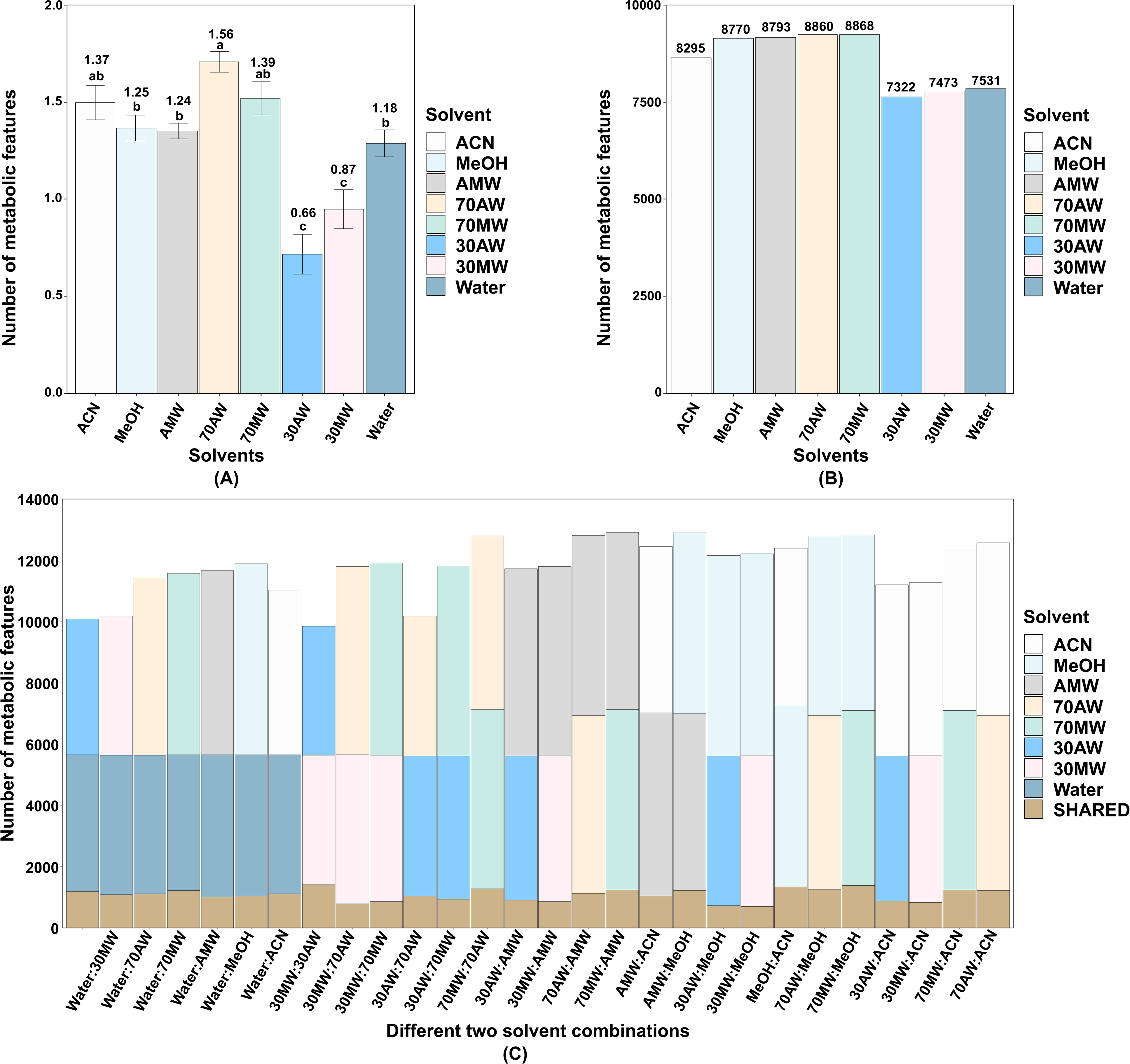
(A) Percentage extraction yield. Comparison of the extraction yield for extracts prepared from 8 extraction solvents using CHs sediments. Bars show the average percent yield of 4 replicate extractions related to the initial dry weight of the sediments, and the error bars represent the standard error. Letter superscripts indicate statistically different groups at a p-value < 0.01 according to Tukey’s HSD test. The average percent extraction yield of each solvent extract is displayed above the letter superscripts. **(B) Single solvent extraction bar plot,** showing a number of metabolic features extracted by eight solvents. Each bar represents a solvent and is displayed with a unique color. The height of each bar indicates the total number of features detected; the exact number is shown at the top of each bar. **(C) Different two parallel single-solvent extract combinations.** Unique features detected by a single solvent from each combination are displayed by each color bar. The common features detected in the two solvent systems are indicated in the grey shade and named ‘SHARED’ features.

### 3.2 Stability, accuracy, and analytical reproducibility of the LC-MS system

A detailed analysis of quality control (QC) samples was performed using the RawHummus tool to evaluate instrument stability, reproducibility, and accuracy. An overlaid TIC (Total ion chromatogram) analysis revealed the least retention time and intensity fluctuations throughout the run (Fig. S6 A). The summed TIC analysis (i.e., the sum of the intensities of all the scan points across the TIC) revealed that global ion intensity variations among the QC samples were within an acceptable range (Fig. S6 B). The TIC correlation analysis revealed that the QC samples had good metabolic profile similarity (Pearson correlation coefficient R >0.90; acceptable range R >0.85) in RT (retention time) and chromatogram peak shape (Table S9), showing the least variance from run to run.

Throughout the chromatogram, RawHummus selected seven putatively annotated ion characteristics (two manually and five automatically) to monitor and assess the LC-MS system for RT, mass, and ion intensity variations. Maximum RT, mass difference, ion intensity ratio, and intensity coefficient of variation (CV) of the extracted ion chromatogram (EIC) of selected seven ion features are presented in Fig. S7 A and B. The maximum RT variation (which should not exceed 1 min) was less than 0.12 min, showing high retention time reproducibility. The mass difference between 0.25 and 1.34 ppm indicated good mass accuracy (maximum acceptable mass variation is up to 5 ppm). The ion intensity ratio (which should be between 1 and 2) was less than 1.6, indicating good stability. The ion intensity coefficient of variation for all m/z values was less than 30%, indicating better instrument stability. One of the m/z features among the seven was 148.0602, which was identified and confirmed as L-glutamic acid after matching its MS/MS spectra with the authentic standard (Fig. S7 C and D).

### 3.3 Detection of features and evaluation of solvent-wise metabolite extraction

A broad range of metabolic features was detected using different solvent combinations. Table 1 depicts the metabolic features discovered and annotated for each extract in both positive and negative modes during early data processing. In RP, metabolic features ranged from the 4898 to 6479 in the ESI (+ve) mode and 2381 to 2759 in the ESI (-ve) mode. Extracts containing organic or organic-rich solvents, such as methanol and acetonitrile combinations, were shown to have more detectable features. The 70MW and 70AW extracts showed the highest detected metabolic features (8868 and 8860, respectively). The order of extracts in terms of total number of the detected metabolic features was 70MW > 70AW > AMW > MeOH > ACN > Water > 30MW > 30AW. The putative metabolites annotations out of total detected features using the KEGG database were between 23.45 and 23.87% in the ESI (+ve) mode and 18.92 to 19.25% in the ESI (-ve) mode. In the HILIC phase, metabolic features ranged from 793-1033 in the ESI (+ve) mode and 2339-3275 in the ESI (-ve) mode. 75HILIC extract showed the highest number of metabolic features detected and annotated, and it was used to compare and combine with RP in subsequent data analysis. The percentage of putative metabolites annotations using the KEGG database was between 28.84-29.82% in the ESI (+ve) and 33.37-33.91% in the ESI (-ve) mode. A large proportion (>75% in RP-LC-MS and >65% in HILIC-LC-MS) of the detected features remained unknown after the database match.

**Table 1:**
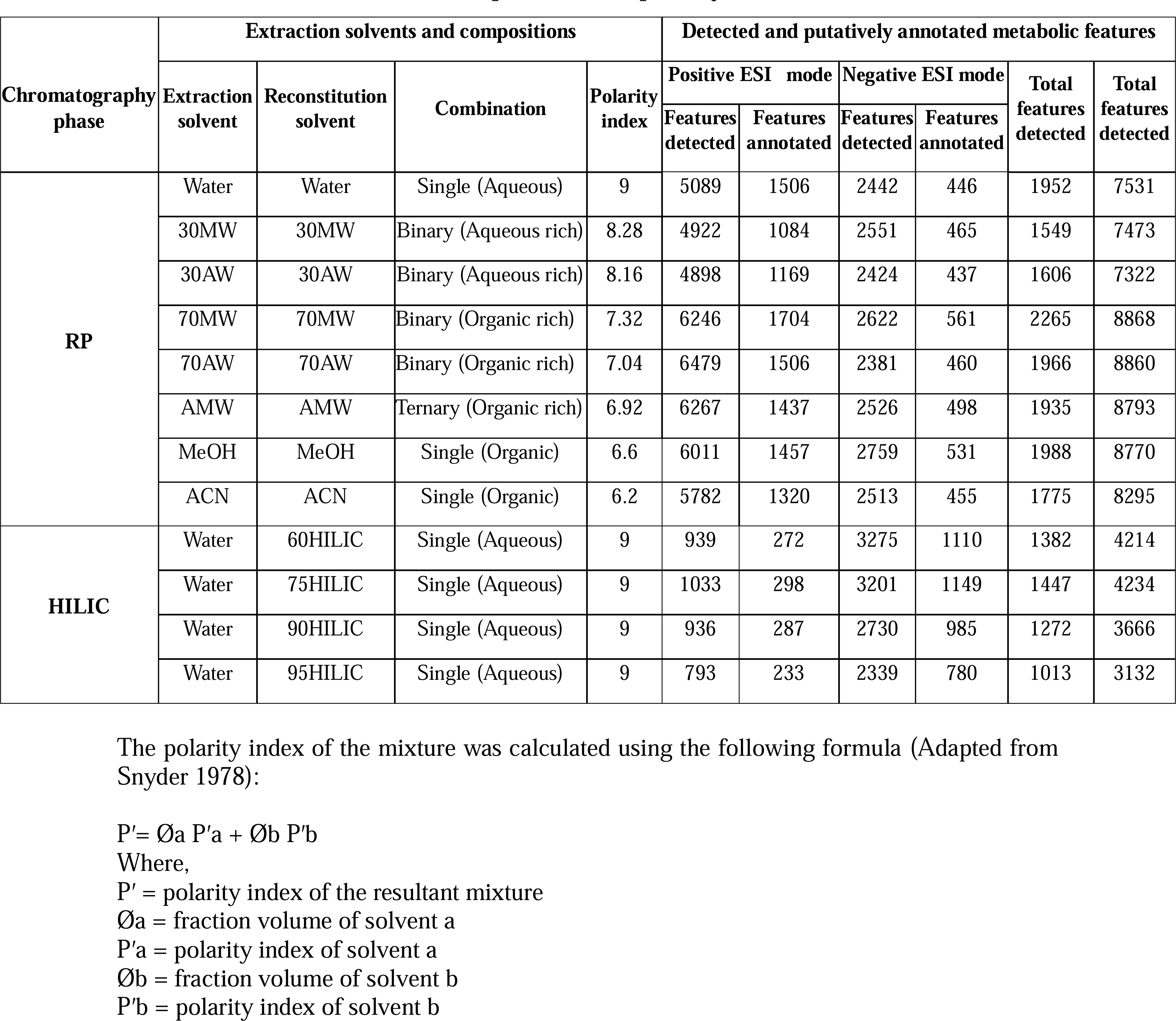
Extraction solvent compositions and polarity index and detected features.

This study aimed to identify the best solvent system, i.e., a single or combined solvent system, that would approach the maximum chemical diversity of CHs. In a single solvent system (Fig. 1B), 70MW solvent extracted the highest number i.e., 8868 of metabolic features. Examining the features from any separately prepared parallel single solvent extractions enables comparing shared and unique features between two or more solvent systems. Shared and unique features in the two- and three-solvent combination are shown in Fig. 1C and Fig. S8, respectively.

### 3.4 Efficiency of extraction and detection of low-abundance metabolic features

To evaluate the sensitivity and dynamic range of our untargeted approach, we examined the proportion of each metabolic feature that contributed to the total signal of metabolic features extracted by the solvents and detected by LC-MS techniques (Ladd et al., 2019). Fig. 2A showed that of the eight distinct solvents, water detected the highest number of low abundant features (267), and MeOH detected the lowest (64), accounting for 50% of the total signal, respectively. Of the four LC-MS techniques, RP in (+)ve mode detected the highest (177), and HILIC in (+)ve mode detected the lowest (8) low abundant features, accounting for 50% of the total signal (Fig. 2B). Notably, RP detected more low abundant features than HILIC, consistent with its more favorable retention and ionization conditions, resulting in enhanced MS detection sensitivity.

**Figure 2:**
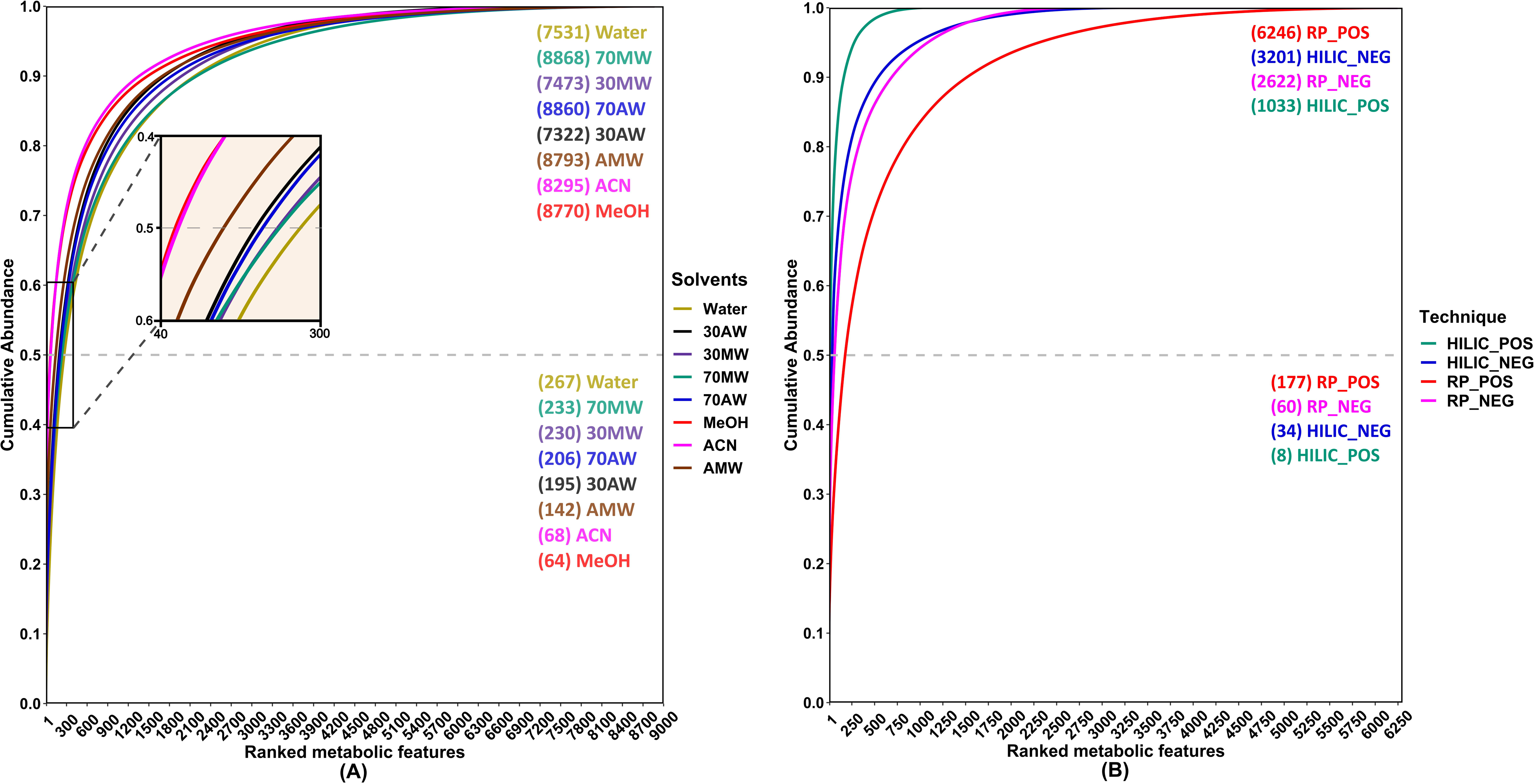
Cumulative abundance plot; for (A) individual solvents, high-quality features are ranked by abundance: 1=most abundant and 2300=least abundant in the case of individual solvents**; for (B) chromatographic phase,** 1=most abundant and 1400=least abundant in the case of the chromatographic phase. The number of LMW DOM features detected by each solvent system and each LC-MS condition accounts for half, and the total cumulative abundance is reported.

### 3.5 Effect of extraction solvents on chemical diversity of metabolites in DOM of CHs

A principal component analysis (PCA, Fig. 3) was performed to understand how solvent extracts were related. The study revealed a tight primary clustering among four biological replicates of each extraction solvent, indicating high analytical reproducibility of the RP-LC-MS technique. PC1 and PC2 accounted for more than 80% variation for all extracts. Notably, the PC1 component alone accounted for 70.4% variation, forming three groups: group 1 (aqueous and aqueous-rich solvents such as water, 30MW, and 30AW), group 2 (organic-rich solvents such as 70MW, 70AW, and AMW), group 3 (organic solvents such as MeOH and ACN). Therefore, secondary grouping, i.e., three distinct clusters, was generated using various extraction solvents. Interestingly, extract clusters were formed away from the PCA plot origin, revealing a particular set of metabolite-enriched clusters.

**Figure 3:**
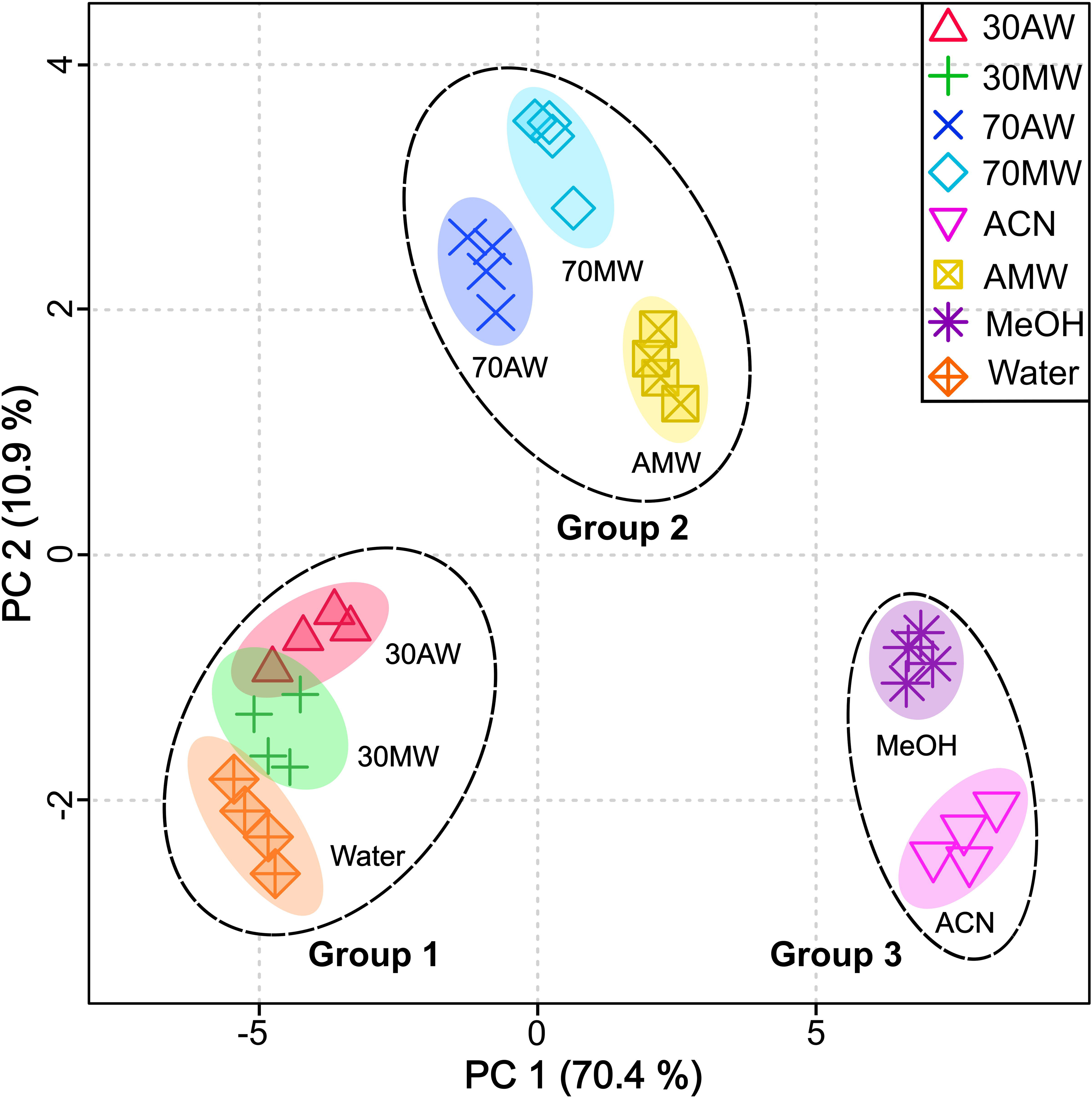
Principal component analysis of eight solvent extracts. PC1 and PC2 accounted for around 80% of the variation in this sample set. Tight primary clusters are produced by replicates of a single solvent system, indicating that each has high reproducibility. Secondary clustering, i.e., groups 1, 2, and 3, are formed by extracts showing the variability of the extracted metabolic feature by respective solvents.

A heatmap (Fig. 4A) analysis shows an abundance of the metabolites of different polarity (polar, mid-polar, and relatively non-polar) and chemical classes based on the polarity of the extraction solvent, from pure aqueous (polar) to pure organic (relatively non-polar) from right to left. The figure shows that using water with no or up to 30% organic solvents (water, 30MW, and 30AW) extracted highly abundant polar metabolites. For example, pyridoxal, 5-oxoproline, L-glutamate, etc., were more prominent in aqueous and aqueous-rich solvents than in the rest of the solvent systems. As the proportion of the organic solvent increased (70MW, 70AW, and AMW), a broad spectrum of polar, mid-polar, and relatively non-polar metabolites (succinate, ectoine, and sphingosine, etc.) were extracted in higher abundance. Using pure organic solvents like MeOH and ACN extracted highly abundant, relatively non-polar metabolites like ecgonine, octadecanamide, dodecanamide, etc. The line plots on the left side of the heatmap showed the pattern of solvent-based metabolite extraction. The magenta lines (Fig. 4B) represent a group of metabolites with high intensities from the top left portion of the heatmap, gradually decreasing from organic and organic-rich to aqueous-rich and aqueous extraction solvents. In contrast, the plot of the green lines (Fig. 4C) showed the opposite trend.

**Figure 4:**
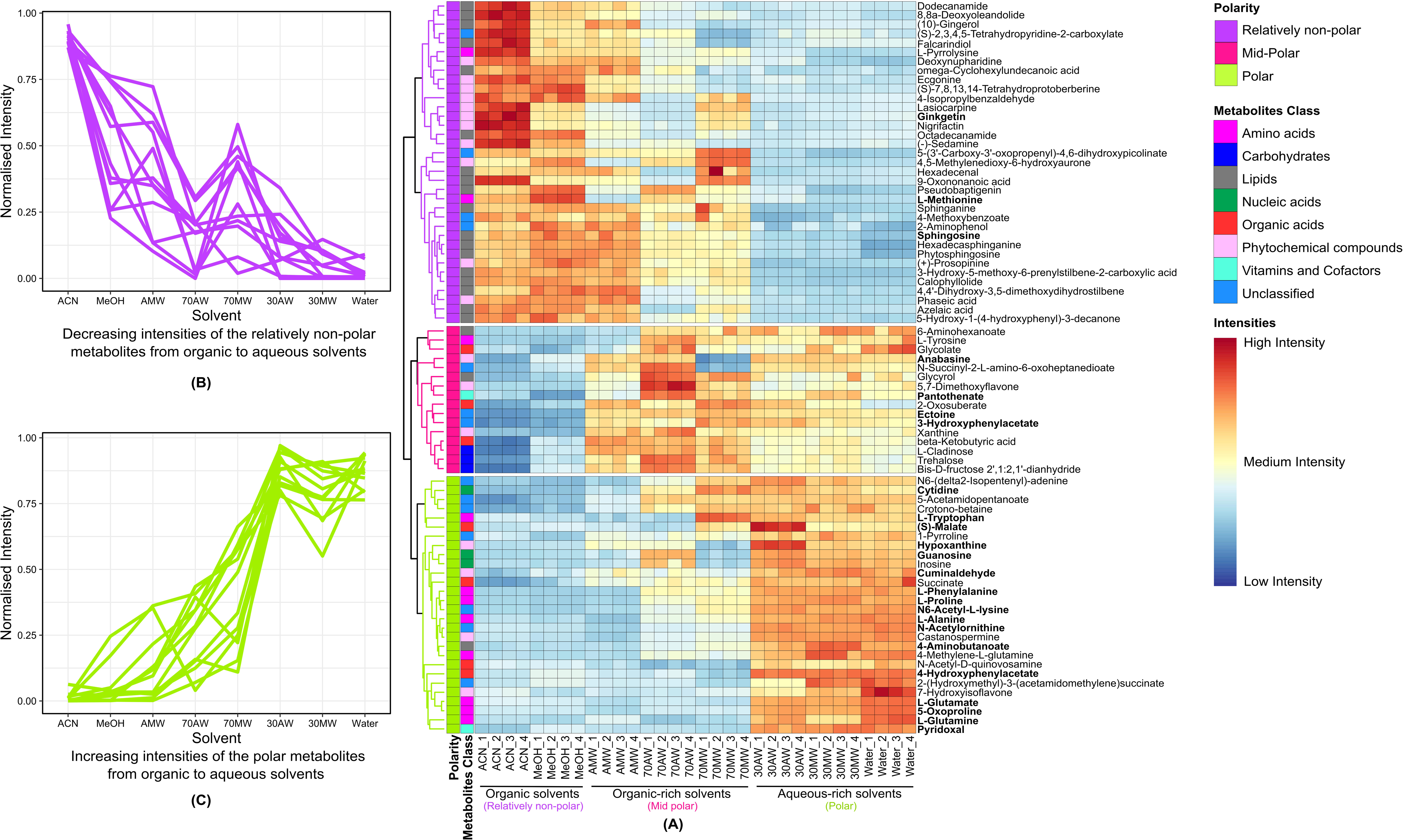
Comparison of extraction conditions and chemical diversity of detected metabolic features. The dendrogram on the left of the heat map shows hierarchical clustering of similarly extracted metabolic features. The heat map displays the intensity of metabolites normalized to the most intense peak within each row (metabolic feature). Differences in metabolic feature peak intensity across extraction conditions are visible as top to bottom – based on polarity and left to right – based on solvents (from organic to aqueous). The line plot at the far left of the figure shows decreasing (magenta lines) and increasing (green lines) peak intensities in extraction conditions from left to right. Metabolic features were roughly divided into polar, midpolar, and relatively nonpolar and categorized into different metabolic classes. Each class is given a different color, as the figure legend shows. Confirmed and putatively confirmed metabolic features after MS/MS spectra match with the standards/predicted theoretical fragments match (via Mass Frontier software) are shown in bold. N = 4 for each extraction condition.

### 3.6 Combination of HILIC and RP chromatography for global metabolomics

The HILIC technique was combined with RP to increase the spectrum of chemically diverse metabolites from CHs. In addition to quantity, qualitative aspects of metabolites are an essential comparison measure. Therefore, to illustrate the distribution of extracted metabolites using both RP and HILIC chromatography, a scatter plot analysis was carried out about retention time (RT) and mass (m/z) dimensions (Fig. S9). In RP, most m/z eluted in the chromatographic gradient’s middle phase (5 to 15 min), while in HILIC, most m/z were eluted at the initial phase (from 0.5 to 10 min) of the gradient.

The Venn analysis and stack bar plot (Fig. 5A, 5B) showed that the best solvent extract from HILIC (75HILIC) with RP best single-solvent (70MW) and two-solvent combination (70MW and AMW), respectively, shared the least number of metabolites in both chromatographic conditions. The HILIC best vs. RP best single-solvent comparison showed that the RP (+ve) mode detected the highest number of unique features (5229), followed by the HILIC (-ve) mode (2457), accounting for 73.68% of the metabolic features across the entire dataset. A similar pattern was observed when HILIC’s best solvent extract was compared with the RP’s best two-solvent combination system. It was noted that most features were detected in a positive mode in the RP and a negative mode in the HILIC technique. Combining HILIC and RP chromatography significantly boosted the total number of metabolic features (Fig. 5C). The number of features when combining HILIC best with the single best RP solvent and best two solvent combinations was 10432 and 16142, respectively. Fig. 5C compares RP and HILIC techniques for detecting highly polar and moderately polar metabolites. The upper row shows high-intensity moderately polar molecules detected in RP, and the bottom row shows high-intensity highly polar molecules detected in the HILIC technique.

**Figure 5:**
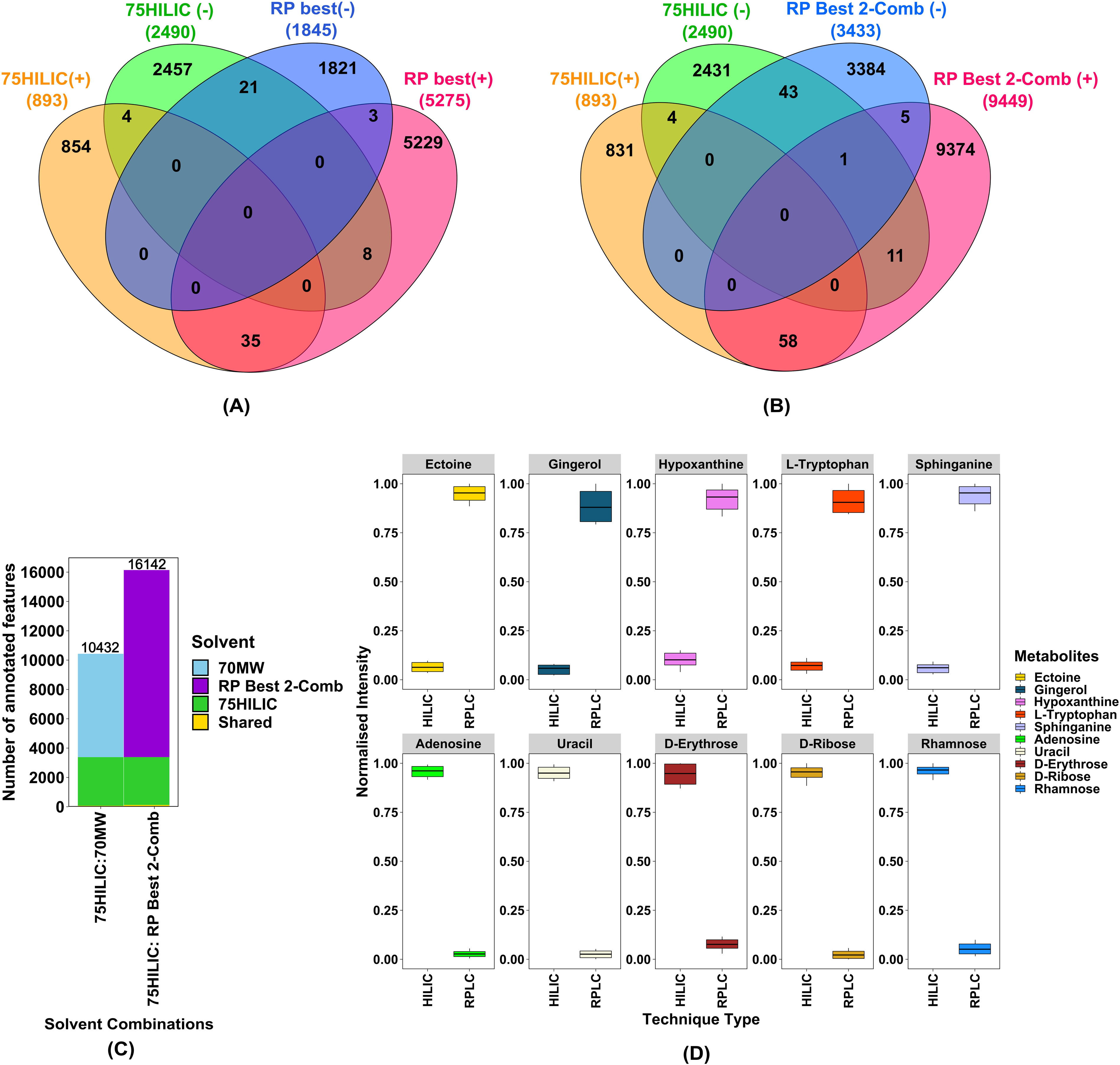
Overlap of metabolic features. detected by HILIC and RP in positive- and negative-ion MS polarities (based on the MS1 neutral mass for the corresponding [M + H] ^+^ or [M − H] ^−^ ion, +/−0.001 Da). **The Venn diagram; (A)** feature detected between HILIC best (75HILIC) and RP best (70MW)**; (B)** feature detected between HILIC best (75HILIC) and RP two best solvent combination (70MW and AMW). **The stacked bar plot (C)** shows unique and shared features between the best HILIC (75HILIC) and RP single solvent (70MW), two solvent combinations. The total number of metabolites detected is displayed on top of each bar. Note that the number of shared metabolites (red) between both techniques is the least in both comparisons. **(D) Intensity comparison of the same metabolites detected by HILIC and RP;** putatively annotated metabolites with high intensity in RP phase (top row) and metabolic features with high intensity in HILIC phase (bottom row). The HILIC phase detected highly polar metabolic features compared to the RP phase, whereas the RP phase detected moderately polar metabolites.

## 4. Discussion

### 4.1 Extraction solvent compositions

In this study, we evaluated several extraction solvents to detect a broad spectrum of metabolites in terms of their number and chemical diversity present in an extractable fraction of the DOM (extra- and intracellular) of CHs. Selecting the best extraction solvent is a pivotal aspect of metabolomics experiments, as efficient extraction relies on the polarity and physicochemical properties of the metabolites and solvent used for extraction (Moosmang et al., 2019). Table S10 lists typical solvents used for the extraction of metabolites from various samples such as human plasma, urine, plant tissue, and soil, including but not limited to water, methanol, acetonitrile, acetone, perchloric acid, ethyl acetate, dichloromethane, chloroform, Isopropanol for analytical platforms like LC-MS, GC-MS, and NMR.

According to Winder et al. (2008), the precipitate formed during extraction with perchloric acid (perchlorate salts) and potassium hydroxide can be problematic if not completely removed, as they are incompatible with analytical platforms like LC and GC. Boiling ethanol caused partial degradation of thermolabile metabolites like pyruvate, nucleotides, and sugars (Villas-Bôas et al., 2005). Acetone is a good choice for extracting polar metabolites, but it showed poor reproducibility among replicates (Bruce et al., 2009). Ethyl acetate, chloroform, 2-propanol, and n-hexane are preferred to extract non-polar metabolites such as lipids (Martin et al., 2014; Swenson and Northen, 2019; Moosmang et al., 2019; Navarrete-Carriola et al., 2024). The solvent that can extract a broad range of metabolites should be less toxic, have high solubility power, good pH tolerance, and be compatible with the analytical platform. Many of these properties can be controlled or modified using appropriate options, e.g., toxicity can be controlled by practicing routine safety measures, and pH can be modified using approved buffers (Mushtaq et al., 2014). However, some properties, like polarity and solubility power, are inherent characteristics of each solvent and can affect the extraction process.

For our study, we selected the most commonly and widely used, less toxic, and highly RPLC-MS-compatible aqueous and organic solvents, such as water, methanol, and acetonitrile. Both methanol and acetonitrile have good miscibility with water and produce homogenous monophasic solutions. Their good wetting behavior allows rapid mixing with the sample matrix. A study by Snyder (1978) (Table S11) classified various solvents according to their polarity index and selectivity group. According to this study, water (polarity index = 9) belongs to the selectivity group of VII and is the most polar solvent. Methanol (polarity index = 6.6), an aliphatic alcohol, belongs to group II and is a protic mid-polar solvent. Acetonitrile (polarity index = 6.2), a nitrile, belongs to group VIa and is an aprotic polar solvent considered relatively non-polar. Also, water has the highest solubility power (>>1), followed by methanol (0.95) and acetonitrile (0.65). Accordingly, we prepared a set of eight solvent systems (Table 1) containing single, binary, and ternary mixtures of these solvents, which can produce solvents of decreasing polarity strength from most polar to relatively non-polar, such as Water > 30MW > 30AW >70MW > 70AW > AMW > MeOH > ACN. Each solvent was evaluated for extraction yield, reproducibility, the efficiency of detecting low abundant features, and the ability to detect the maximum number and chemically diverse set of metabolites.

### 4.2 Metabolites extraction using appropriate extraction solvents and their combinations

High throughput techniques like LC-MS-based untargeted metabolomics can detect thousands of metabolites to characterize the nearly complete metabolome. This allows assessment of the total feature count, i.e., the total number of metabolite features detected as an important parameter to compare the influence of the extraction solvents extracting the highest metabolite number (Hemmer et al., 2024). Nevertheless, the solvent that extracts maximum metabolite features is not always the one that provides the broadest metabolome coverage due to artefactual interferences such as contaminations during metabolite extractions, column carryover from previous experiments, background noises detected by MS, and misannotation of the data during bioinformatic processing (Mahieu et al., 2014). The same sample/matrix can be extracted with different solvent choices as per the study requirement. For example, for the fecal metabolite extraction, Huang et al. (2013) used MeOH, Jiménez-Girón et al. (2015) used saline water, and Deda et al. (2016) used aqueous ACN.

The choice of the extraction solvent could be study-specific to meet the desired objectives of the study in question. Methanol and aqueous methanol are the most popular choices as extraction solvents in many studies due to their ability to extract a broad and diverse set of metabolites (like amino acids, organic and nucleic acids, secondary metabolites, and sugars) compared to water and ACN (Cheng et al., 2020). In the current investigation, binary mixtures of organic-rich solvent-mixtures of MeOH and ACN extracted the highest number of metabolites (70MW and 70AW, Fig. 1B). We observed that organic solvents extracted more metabolites than aqueous solvents. However, adding a small fraction of organic solvents in water slightly decreased metabolic features. Meanwhile, adding a high fraction of organic solvents increased the number of metabolic features. Differences in organic and aqueous extract could be due to the more penetrability of cold organic solvents, thereby disrupting microbial cell membranes and resulting in leakage of intracellular contents, increasing overall extraction efficiency (Pinu et al., 2017).

To maximize the number and diversity of metabolic features to be extracted, we considered metabolic features from any separately prepared parallel single-solvent extractions of the eight extracts for comparing shared and unique features with the pairs of two-solvent combinations, as shown by Martin et al. (2014). The total number of features and the shared features differed significantly, depending upon which solvent combination was used. The 70MW: AMW was the most effective extraction solvent pair in two-solvent combinations (Fig. 1C). The heatmap analysis further supported this, showing that 70MW extracted high-intensity metabolites in the region where AMW showed less intense metabolite extraction. This could be due to differences in the polarity of the metabolites extracted. One can select complementary two-solvent combinations of parallel or repeated extractions to increase this number further. However, the combinations of solvent extraction, either parallel or repeated, are well suited to small-scale studies. Implementing large-scale experiments or routine analysis makes the work more time-consuming and laborious and may result in diluted metabolic extract (Moosmang et al., 2019). Moreover, multistep protocols involving repeated extractions will likely introduce relatively higher deviations and variations in metabolite recovery (Karu et al., 2018).

### 4.3 Influence of the extraction solvent on the chemical diversity of the metabolites

Metabolites represent the molecular endpoint of gene expression, and metabolomics offers a comprehensive approach to understanding the organism’s phenotype or habitat (e.g., CHs). For instance, human blood serum comprises more than 20,000 known chemically diverse metabolites, and this number expands to over 200,000 annotated compounds when exogenous metabolites derived from food, microbiota, and drugs are included (Wishart et al., 2022). According to the Yeast Metabolome Database (YMDB), over 16,000 chemically diverse metabolites are present in *Saccharomyces cerevisiae* (Ramirez-Gaona et al., 2016). Sessile organisms like plants display far greater metabolic diversity than other organisms, with the plant kingdom commonly being stated to contain between 200,000 and 1 million metabolites (Dixon and Strack, 2003; Rai et al., 2017; Fang et al., 2019). In respective organisms, these metabolites belong to different chemical classes like acids (amino, organic, and nucleic), sugars, lipids, and phytochemicals (alkaloids, flavonoids, isoprenoids, glycosides, etc.). It is practically impossible to extract all metabolites with any single solvent. Notably, the total number of extractable, detectable, and yet identifiable metabolites is extensive, which requires systematic assessment of extraction solvents to extract the maximum possible metabolites followed by complementary separation, detection, and analytical platforms for unbiased screening of the metabolome (Kind et al., 2009; Kuehnbaum and Britz-McKibbin, 2013).

One of the objectives of our study was to evaluate the ability of various extraction solvents to extract a large number of chemically diverse metabolic features. The polarity of the extraction solvent, whether polar, mid-polar, or non-polar, is a critical factor in extracting these diverse metabolites. The type of sample also influences the chemical nature of the extracted metabolic features. Therefore, we propose that the overall chemical diversity of the extracted metabolites is a combined effect of the extraction solvent’s polarity and the sample type. Moosmang et al. (2019) discussed the scope of using aqueous (water) and organic (methanol and acetonitrile) as extraction solvents for different chemical classes of metabolites. According to their study, water can extract highly polar (hydrophilic) molecules, whereas methanol and acetonitrile mainly cover moderately polar and relatively non-polar molecules of the sample metabolome. The main advantages of water as an extraction solvent are that it is cost-effective and reproducible; the extract can be directly used for LC-MS with minimal further treatment and is non-toxic. In our study, to increase chemical diversity from hydrophilic to lipophilic, we prepared extraction solvents ranging from water (aqueous, highly polar) to methanol (organic, mid-polar) and acetonitrile (organic, relatively non-polar) and their combinations (aqueous, and organic).

The heatmap analysis (Fig. 4) depicted the influence of extraction solvents on the abundance of extracted metabolites and their clustering in different chemical classes and polarity. From the figure, it can be concluded that adding the organic solvents changed the polarity of the solvent and, accordingly, the extraction of the metabolites. Consequently, a broad range of chemically diverse metabolites could be extracted. For particular questions like targeted metabolomics, pure organic solvents like acetonitrile or methanol could extract lipophilic fractions, while water could extract highly hydrophilic fractions.

### 4.4 Combining HILIC and RP as an excellent approach to increase metabolome coverage

To this point, we have discussed how the meticulous selection of an extraction solvent can enhance the overall number and chemical diversity of the metabolites. At the same time, careful selection of an appropriate analytical platform is crucial for identifying a large number of metabolites. With high-throughput technology and software advancements in the last couple of decades, LC-MS has emerged as the top choice as an analytical platform for metabolomics and has been extensively used for many studies involving targeted or untargeted metabolomics of the plant and human tissues, urine, and bacterial cultures (Moosmang et al., 2019; Schippers et al., 2023; Navarrete-Carriola et al., 2024). RP-LCMS stands out among all analytical platforms for its robust, high-throughput, and reproducibility for scientific findings. It is especially adept at detecting mid-polar to non-polar molecules (Hemmer et al., 2024). However, highly polar and ionic molecules (e.g., amino, organic acids, and sugars) are often poorly retained and resolved on RP-LC columns, hindering their detection and accurate quantification (Bieber et al., 2017). For a complex mixture such as soil, sediment, and similar environmental samples where polar molecules (e.g., water-soluble fraction of the DOM of CHs ecosystems) constitute a significant portion of the metabolome, it is advised to combine RP-LCMS with the HILIC technique, the latter being an excellent analytical platform to retain and detect highly polar molecules, thus complementing RP-LCMS (Ladd et al., 2021).

In our study, we used dual LC-MS (RP and HILIC) and dual ionization mode (positive and negative) approaches to increase the coverage of the metabolome in Antarctic CHs. Qualitative characteristics of the metabolic features regarding their distribution in chromatographic and mass spectral dimensions (Fig. S9) varied in RPLC and HILIC. In RPLC, the feature distribution of extracts was heavily weighed between RT 5 min and 15 min, indicating retention of relatively polar and non-polar molecules. In contrast, the feature distribution of extracts was denser between 0.5 and 10 min in HILIC, indicating well-retained highly polar molecules. The differential detection of highly polar and relatively non-polar molecules revealed the high intensity of highly polar molecules in the HILIC compared with RPLC, which shows more intense, relatively less polar molecules. To evaluate this, we selected some putatively annotated highly polar and moderately polar metabolites for comparison (Fig. 5D). Moderately polar molecules like gingerol, hypoxanthine, L-tryptophan, etc., were detected at high intensity in RP-LCMS. In contrast, highly polar molecules like D-ribose, uracil, rhamnose, etc., were detected in high intensity in HILIC. This result is consistent with the fact that the HILIC approach is better suited for detecting highly polar molecules. At the same time, RP-LC is better at detecting mid- or relatively non-polar molecules. Analyzing the overlap between the two conditions revealed that the least metabolic features were shared between these two techniques (Fig. 5A and 5B). Only about 0.68% of metabolic features were shared in more than one technique in a single solvent and two-solvent systems, respectively. Contrepois et al. (2015) showed that combining the HILIC and RPLC-MS approaches greatly expanded metabolome coverage, with 44% and 108% new metabolic features detected compared to RPLC-MS alone for urine and plasma samples, respectively.

Similarly, in the present study, combining HILIC with RP-LC for a single, two-solvent combination led to the detection of 46.96% and 24.52% of unique metabolic features, respectively, compared to RP-LC alone (Fig. 5C). The decrease in the detection of unique metabolites by combining more than one solvent was because the contribution of the HILIC remained the same; while using RP increased the number of metabolites in a two-solvent combination. Hence, to approach maximum coverage and chemical diversity of the LMW metabolites present in the DOM of CHs, using both (RP and HILIC) chromatographic techniques should be considered whenever possible. Employing dual-LC and dual ionization polarity, which has not been reported to characterize the range of the LMW of DOM from CHs, adds a unique perspective to our study.

### 4.5 Detection of low abundance metabolites, reproducibility, and extraction yield

An untargeted metabolomics approach should reliably detect a diverse group of compounds, including both high- and low-abundant metabolites. The climate in Antarctica is severe due to scarce nutrient availability and extremely low temperatures, which may result in conditions where metabolic traits in the DOM with low abundance could also be biologically significant. For arctic and forest soils, it was observed that microbes would prefer low-concentration compounds and cycle through the soils faster, contributing disproportionately to the DOM fraction that is mineralized into CO_2_ and CH_4_ (Van Hees et al., 2005; Drake et al., 2015; Ladd et al., 2019). In Antarctic CHs, liquid water is available briefly during austral summer from perennial ice meltdown (Luis et al., 2022). It could provide water-soluble nutrients or metabolites to microbes for their growth and metabolism. Probably because of this, water detected the highest low-abundant features in the CHs DOM (Fig. 2 A). This suggests that the inhabiting microbes prefer water-soluble metabolites for their metabolism. Thus, water, a highly polar solvent, can be a good starting point for characterizing the metabolomes of these environmental ecosystems. In the chromatographic technique, RP detected more low-abundant features than HILIC, which could be due to more favorable retention and ionization conditions, leading to enhanced MS detection sensitivity (Fig. 2B).

Using unique metabolic features and corresponding normalized peak areas, we evaluated the reproducibility of the solvents for the untargeted measurement across replicates of the eight solvent systems using principal component analysis (PCA) (Fig. 3). All extracts produced consistent overall reproducibility. The tight primary clustering formed among the replicates of each solvent showed good analytical reproducibility of the respectively extracted features under varied analytical conditions. For metabolic features reproducibility, we assumed a coefficient of variation (CV) of < 25% among the four replicates of each extraction solvent in positive and negative ionization modes for RP and HILIC techniques (Fig. 6A). The percentage of the filtered features was higher in organic solvents among other extractions in RP and was highest in 75HILIC (reconstituted with 75% ACN: 25% water) among other HILIC extracts. The mean of all features within extraction solvents (Fig. 6B) shows the majority of RP and HILIC extracts were close to or below the set limit (< 25%) of the CV. In HILIC, the extract is reconstituted with 60% ACN (60HILIC, ran in positive mode), and the extract reconstituted with 90% ACN (90HILIC, ran in the negative mode) showed the highest mean CV and, thus, poor reproducibility. According to Manier and Mayer (2020), solvents used to reconstitute dried extract in HILIC have an impact on recovery and, therefore, result in the detection of the features with varying consistency. Hence, they suggested optimizing the reconstitution solvent for every chromatographical system used, irrespective of the chemical classes of the molecules to be analyzed. Our investigation found that reconstitution with 75% ACN (i.e., 75HILIC) showed better results than other HILIC reconstitutions.

**Figure 6:**
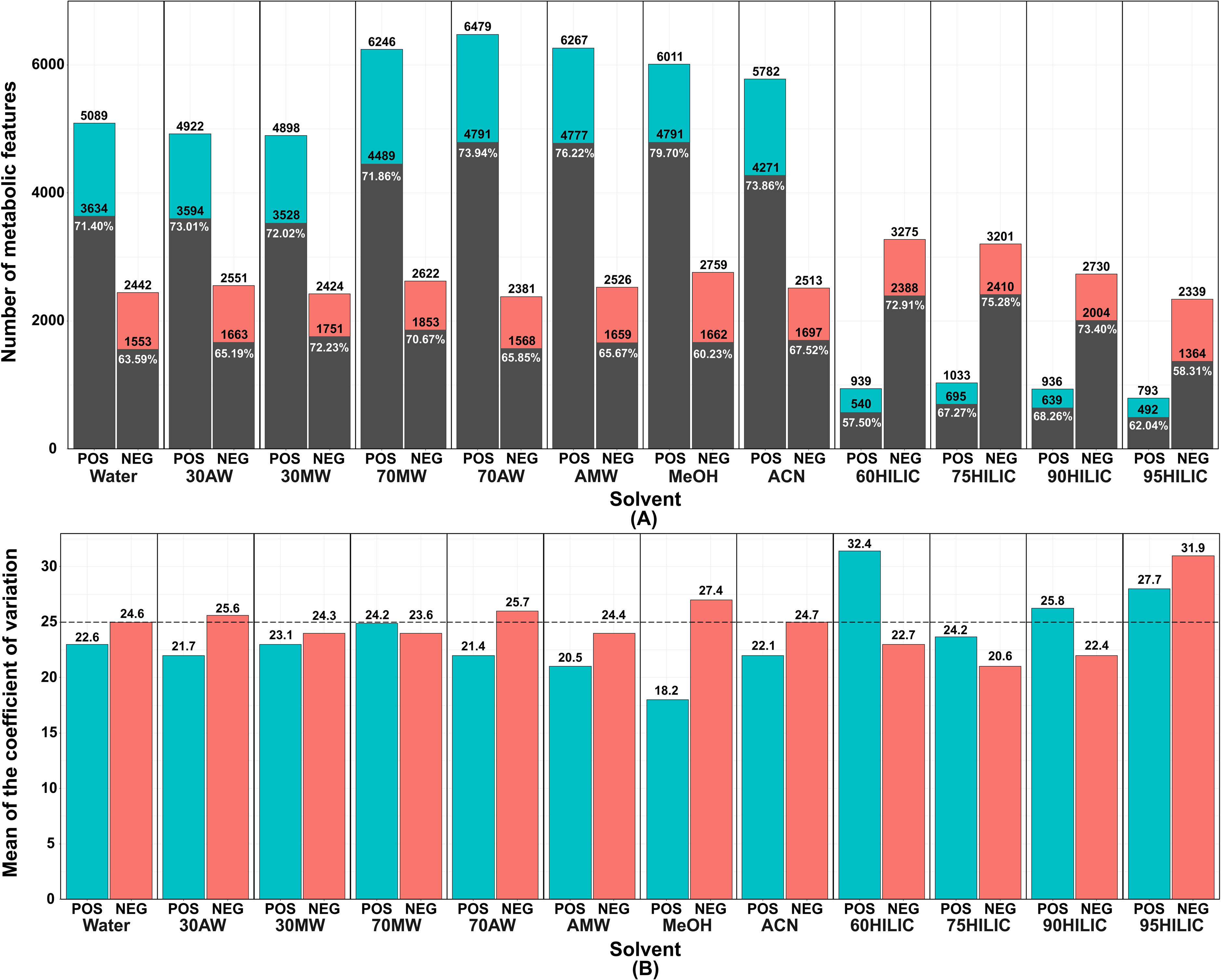
Metabolic features reproducibility; **(A)** The total number of metabolic features detected are shown at the top of each bar in positive (POS) and negative (NEG) ionization mode in RP (first eight solvents from left) and HILIC (last four solvents) technique. The coefficient of variation (CV<25%) applied, and the resulting number of metabolic features are shown at the top of the dark bar. The percentage of these features is shown in white font. **(B)** shows the mean of the coefficient of variation of the total detected features at the top of each bar in positive (POS) and negative (NEG) ionization mode by each RP (first eight solvents from left) and HILIC (last four solvents) technique. The black dashed horizontal line at the y-axis marks the set limit of CV<25%.

Measuring the extraction yield after removing the extraction solvent reflects crude extraction efficiency in quantity. However, it may not indicate the exact chemical diversity of the extract. Also, the colors of the extract (dark, faint, or colorless) do not correlate with the extraction efficiencies as it could be due to co-extraction of macromolecules such as chlorophyll pigments and phycobiliproteins present in cyanobacteria and microalgae (Pagels et al., 2021). In terrestrial soil samples, co-extraction of humic acids imparts a dark color to the extract and may interfere with detecting other biologically significant molecules, e.g., DNA or proteins (Lakay et al., 2007; Qian and Hettich, 2017). We found that 70AW and 70MW were the most efficient solvents in extraction yield compared to pure water, methanol, and acetonitrile for extracting metabolites from CH samples. 70MW, despite being a colorless extract (Fig S5 A), produced a high extraction yield. This result shows a similar observation in a previous study where aqueous organic extracts yielded more than absolute organic extracts, and colorless extract also resulted in a high extraction yield (Sultana et al., 2009; Martin et al., 2014). In conclusion, a high-yielding solvent may produce extracts of low overall quality due to limited chemical diversity.

### 4.6 Roles of the detected metabolites

In the present study, CHs sediment LC-MS-based untargeted metabolomics workflow detected a wide range of small molecules such as carbohydrates, acids (amino, nucleic, and organic), lipids (fatty acyls, glycerolipids, glycolipids, etc.), phytochemicals (alkaloids, flavonoids, and isoprenoids, etc.), and vitamins using RP and HILIC techniques. In recently published studies, the authors applied metabolomics to reveal fundamental metabolic processes essential for sustaining life in cold environments. For example, a study by Gokul et al. (2023) on cryoconite samples from Foxfonna ice caps in the Arctic found higher levels of TCA cycle intermediates like fumarate, succinate, and isocitrate, indicating high ATP generation, which could be linked to higher microbial activities like bioenergetic metabolism with carbohydrates, amino acids, and fatty acid degradation. This study also indicated that microbes in such nutrient-deficient conditions prefer energetically cheaper degradative pathways to generate metabolites to sustain cellular growth, particularly concerning nitrogen-containing molecules. It is known that microbial growth is mediated more by nitrogen than soil carbon content (Eisenhauer et al., 2010). This indicates that amino acids are crucial in regulating these microbial communities.

Vitamins like riboflavin, produced by bacteria, were found to promote freshwater microalgal cell growth by mitigating salt stress (Palacios et al., 2021). The presence of transcriptional regulator AZeR is mainly limited to the Proteobacteria phylum (Shibl et al., 2020), and the bacterial response to azelaic acid is controlled by AZeR (Bez et al., 2020). Doting et al. (2023) detected riboflavin and azelaic acid in cryoconite samples from Greenland ice sheets and suggested further research into the ability of microbes habituating these surface ice to produce or sense these molecules for a better understanding of microbial interactions. Production of small osmoprotectants and cryoprotectants like glycine betaine aldehyde, glycerol, trehalose, sucrose, and D-sorbitol can allow cryoconite microbial community to reduce the freezing point of cytoplasm thereby preventing macromolecule aggregation and can also stabilize cell membrane under cold conditions (Tribelli and Lopez, 2018). Membrane lipid unsaturation with branched and unsaturated fatty acids by cell membrane modifications is a crucial response to cold stress by the habituating microbes (Siliakus et al., 2017; Edwards et al., 2020). Therefore, in cold and challenging environments like CHs, microbes might have found a way to survive and proliferate by developing adaptation strategies to adapt to their surroundings. This supports the detection of amino acids and carbohydrates, intermediates of the TCA cycle, vitamins like riboflavin, lipids, and molecules like azelaic acid (Table S12) in the current investigation, which may help to better understand their roles in adaptation strategies by microbes in icy environments such as CHs. It is important to note that our aim in this study was not to identify each feature detected but to evaluate extraction solvents for an untargeted approach to reveal information-rich molecular profiling of LWM DOM present in the unique and complex matrix like CHs. Further confirmation of features using appropriate metabolic standards is required to increase the reliability of future metabolomics studies.

## 5. Conclusions

This study comprehensively assessed a defined set of solvents for an RP-LCMS-based high-throughput, sensitive, and robust, untargeted metabolomics workflow. This method can be feasibly applied to extract a broad spectrum of the LMW molecules in the DOM of global CHs. Aqueous solvents like water extracted highly polar and low-abundant metabolites, while pure organic solvents like methanol and acetonitrile extracted more non-polar metabolites. The organic-rich solvent mixtures (70MW and 70AW) led to the extraction of a large number and broad range of chemically diverse metabolites. All solvents showed good reproducibility. No solvent could extract a complete feature set, so combining parallel single or repeated extractions with multiple solvent systems is required to enhance the number and capture maximum chemical diversity. A combination of HILIC and RP chromatography expanded the coverage of the diverse and large-scale chemical features, providing qualitative and relative quantitative data and yielding an information-rich chemical snapshot of biogeochemical activity in cryoconite holes. We conclude that the choice of the solvent and technique is exclusively study-specific, and solvents can be preferred as per the requirement of the experiment.

Our research opens up exciting possibilities for future studies. We suggest the application of a 13C isotope-assisted workflow that would allow the direct consideration of detectable metabolites of biological origin to assess different extraction mixtures. This could significantly enhance the accuracy and reliability of future studies. The future work should include applying the technique across multiple northern and southern hemisphere cryoconite hole samples, correlating shifts in LMW DOM chemistry with microbial community composition and environmental variables to map compounds to metabolic pathways. Furthermore, the larger-scale studies integrating metabolomics with other ‘omics’ approaches will provide deep insight into critical biogeochemical transformations, e.g., C and nutrient cycling over time or space. Detailed stoichiometric information for mechanistic models will help reduce uncertainty in predictions of how cryoconite hole ecosystems of the northern and southern hemispheres will respond to climate change.

## 7. Data availability

The high-resolution LC-MS data (MS1 and MS2) from the present study are available at the GNPS-MassIVE repository with MassIVE ID: MSV000094568.

FTP Download Link: ftp://MSV000094568@massive.ucsd.edu

## Acknowledgments

We thank the Director, National Centre for Polar and Ocean Research (NCPOR), Goa, India, for providing access to the Antarctic cryoconite hole samples, Dr. B Santhakumari for providing access to the Mass spectrometry facility at CSIR-National Chemical Laboratory, Pune and Mr. Anirban Majumdar for helping with the R scripts. This work was funded by the Ministry of Earth Sciences (MoES), Government of India, under a central sector umbrella scheme named “Polar Science and Cryosphere Research (PACER).”

## Abbreviations

CHs: Cryoconite holes
DOM: Dissolved organic matter
EIC: Extracted ion chromatogram
ESI: Electrospray ionization
FTICR-MS: Fourier transform ion cyclotron resonance-mass spectrometry
GC-MS: Gas chromatography-mass spectrometry
HESI: Heated electrospray ionization
HILIC-LCMS: Hydrophilic interaction liquid chromatography-mass spectrometry
HRMS: High-resolution mass spectrometry
LC: Liquid chromatography
LC-MS/MS: Liquid chromatography-tandem mass spectrometry
LMW: Low molecular weight
NMR: Nuclear Magnetic Resonance
PCA: Principal component analysis
QC: Quality control
RP-LCMS: Reverse phase liquid chromatography-mass spectrometry
UHPLC: Ultra high-pressure liquid chromatography
Water: 100% water
30MW: 30:70 Methanol: Water
30AW: 30:70 Acetonitrile: Water
MeOH: Methanol (100%)
ACN: Acetonitrile (100%)
70AW: 70:30 Acetonitrile: Water
70MW: 70:30 Methanol: Water
AMW: 40:40:20 Acetonitrile: Methanol: Water
60HILIC: 60: 40 Acetonitrile: Water
75HILIC: 75: 25 Acetonitrile: Water
90HILIC: 90: 10 Acetonitrile: Water
95HILIC: 95: 5 Acetonitrile: Water

## Workflow 1

The workflow for the untargeted metabolomics approach established to analyze LMW DOM from Cryoconite hole samples. Quadruplets of the extracts for each solvent system were done separately. The resulting concentrated aliquots (samples) were run on two LC phases (RP and HILIC) and in two MS modes, ESI (+ve) and ESI (-ve), resulting in four analytical conditions per sample.

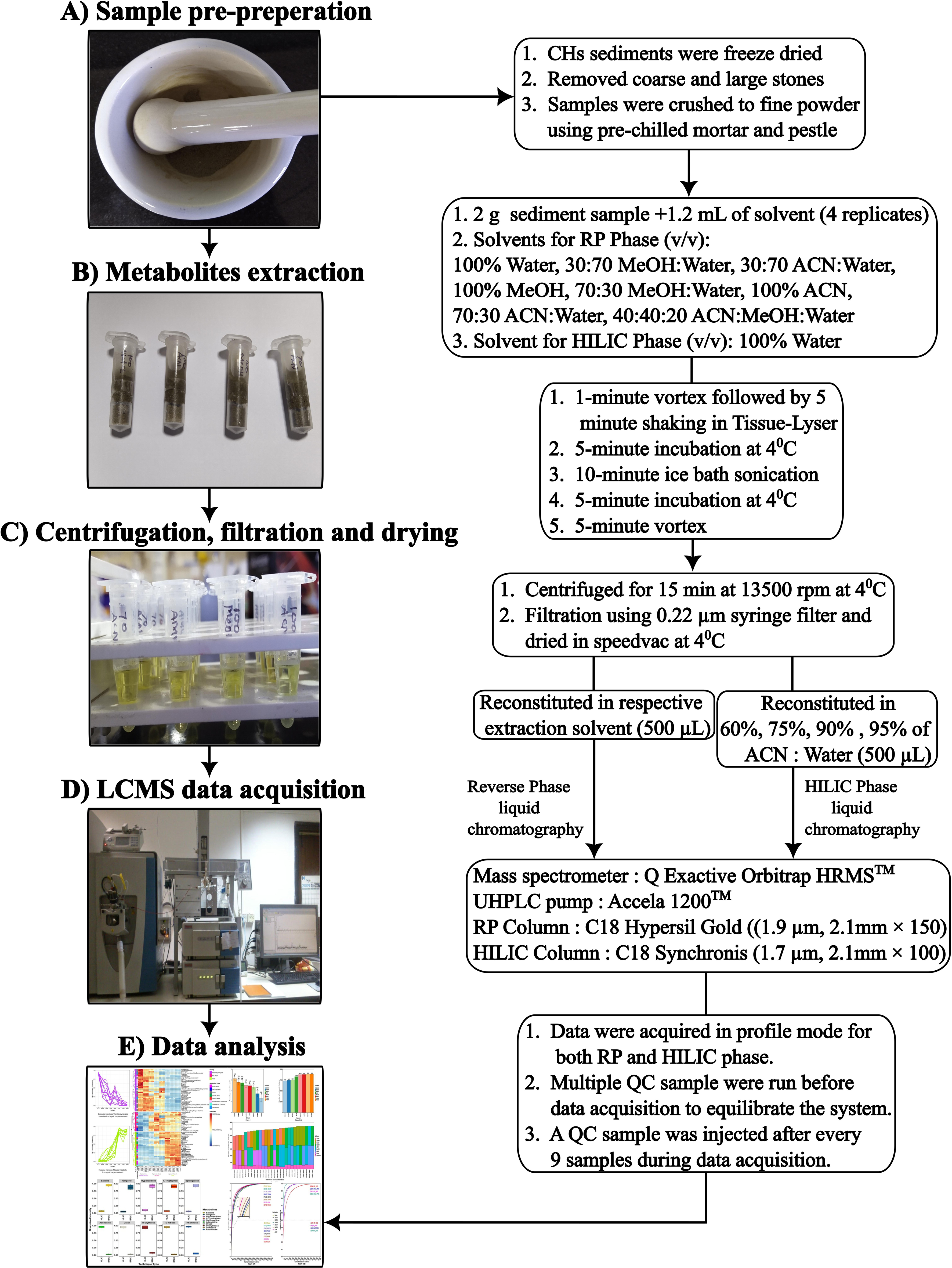

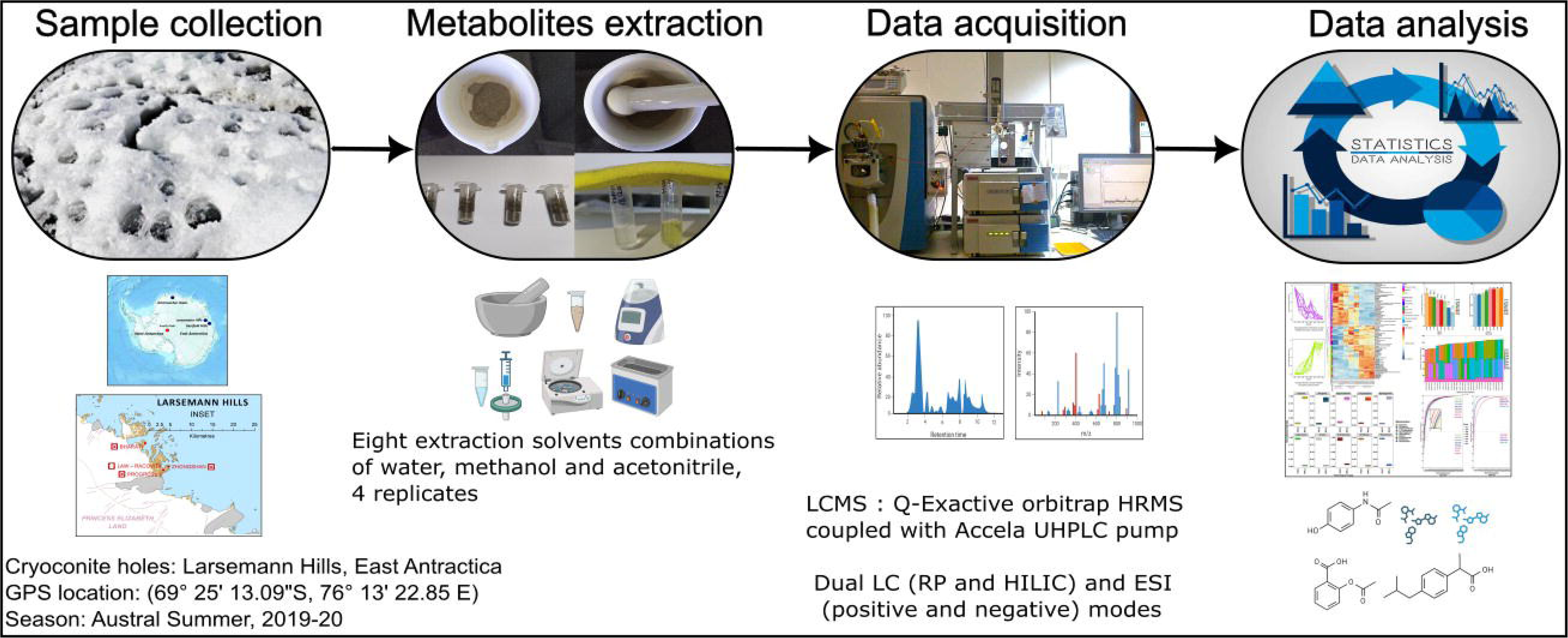

## Highlights

- First report on evaluating extraction solvents for metabolomics of cryoconite holes
- Eight solvents were assessed for the highest and most diverse metabolite extraction
- Organic-rich MeOH: Water (70:30 v/v) solvent extracted the highest metabolites
- Dual LC and ionization modes increased the breadth of the detected metabolites
- Lays foundation for future metabolomics work of global supraglacial habitats

