## Supplementary material for "Evaluation of extraction solvents for untargeted metabolomics to decipher the dissolved organic matter of Antarctic cryoconite holes": Supplimentary_Figures.pdf

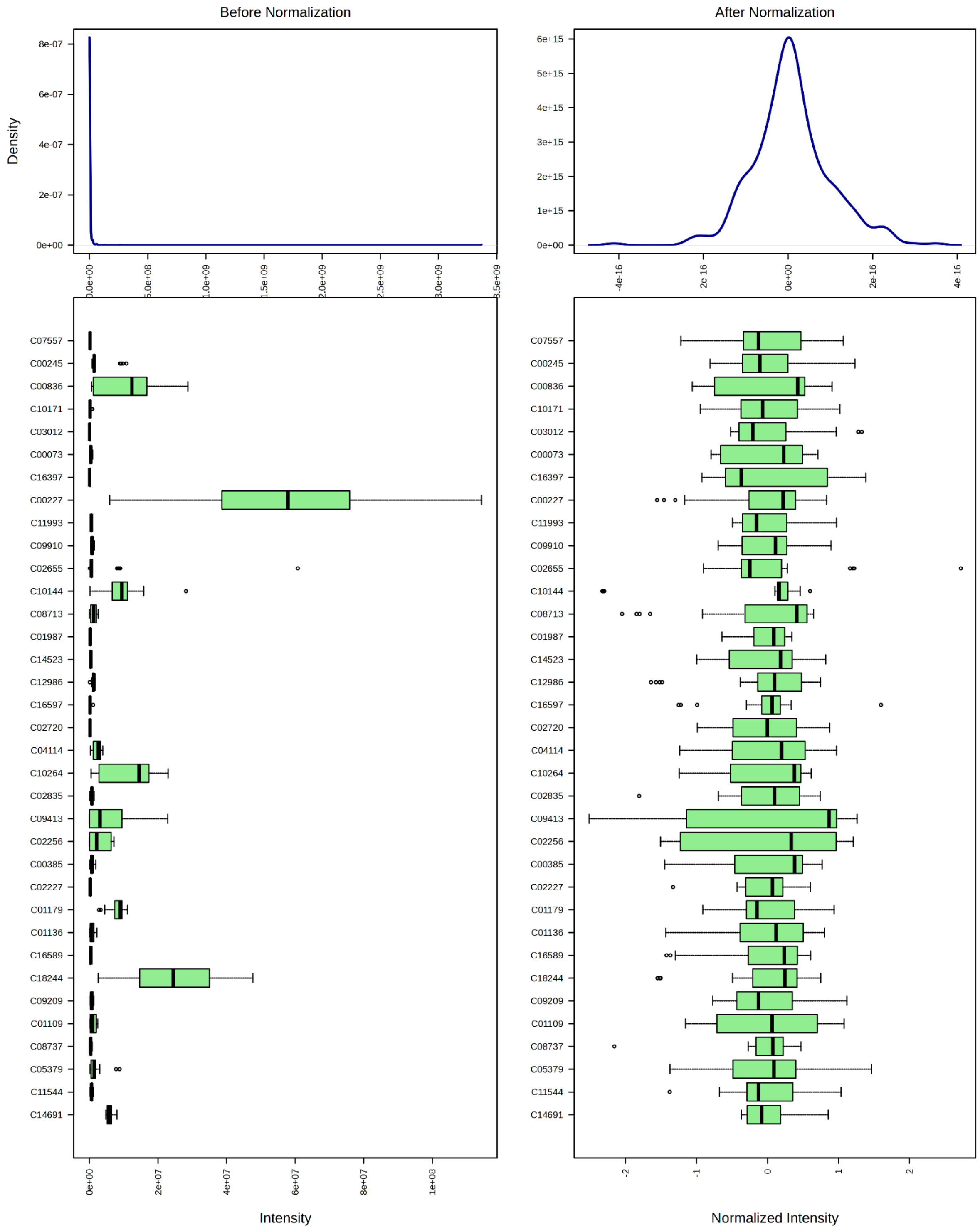


**Fig S1: Distribution of the data before and after normalization**

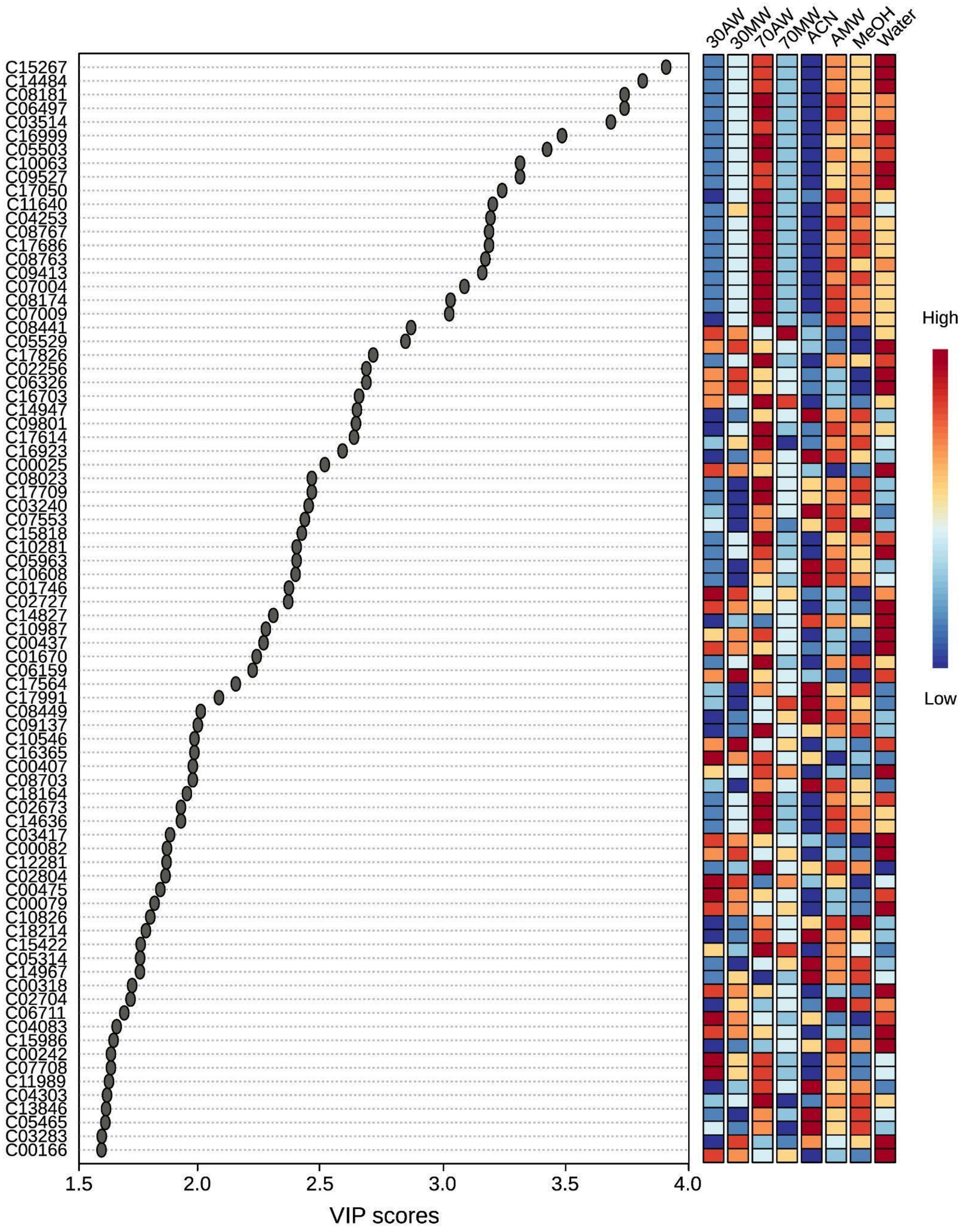


**Figure S2: PLS-DA VIP (variation in projections) features > 1.5**

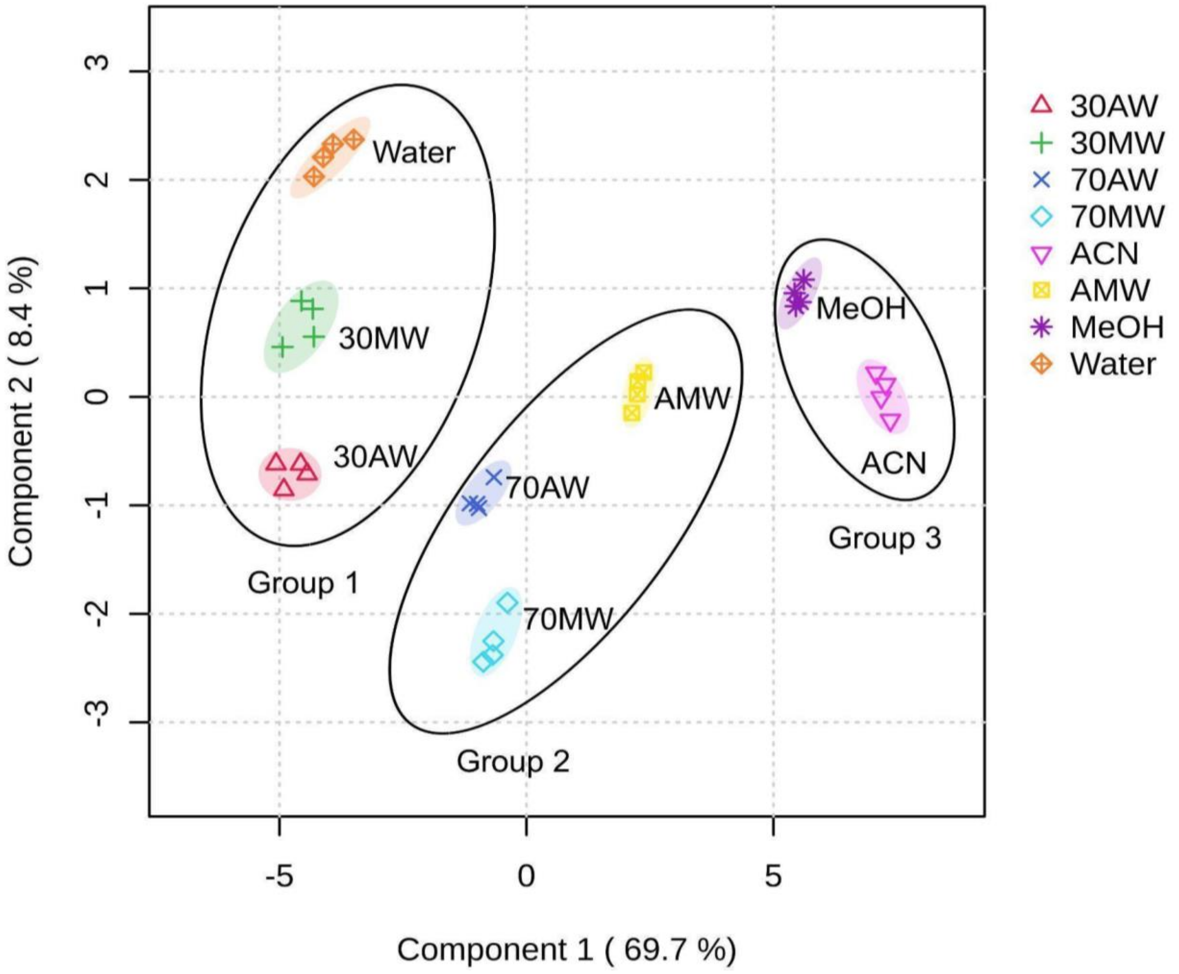


**Figure S3: PLS-DA scores plot**

**Figure S4 (below): Metabolic features validated by MS/MS spectra match with authentic standards**

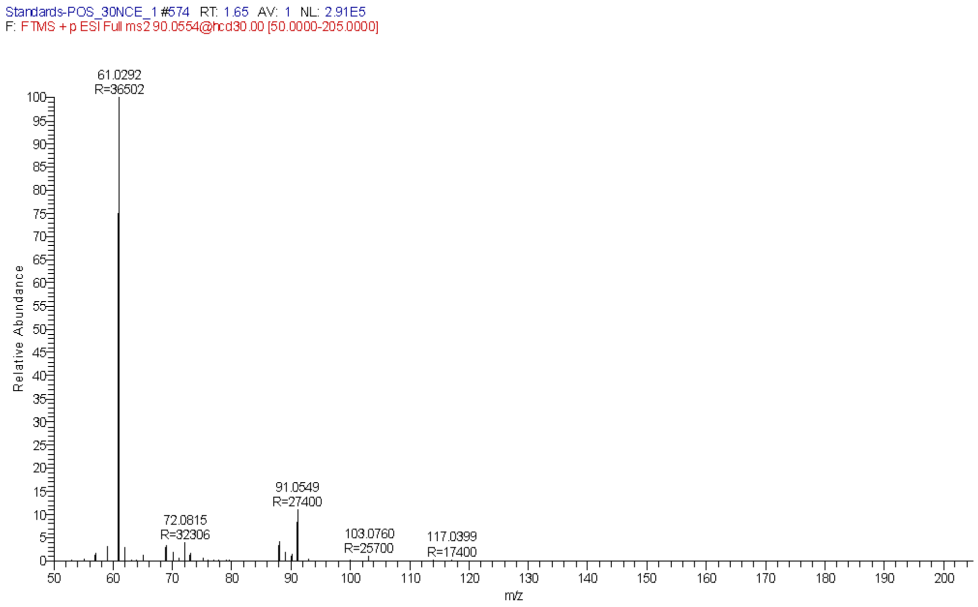


Experimental MS/MS spectra of the authentic standard of L-Alanine.

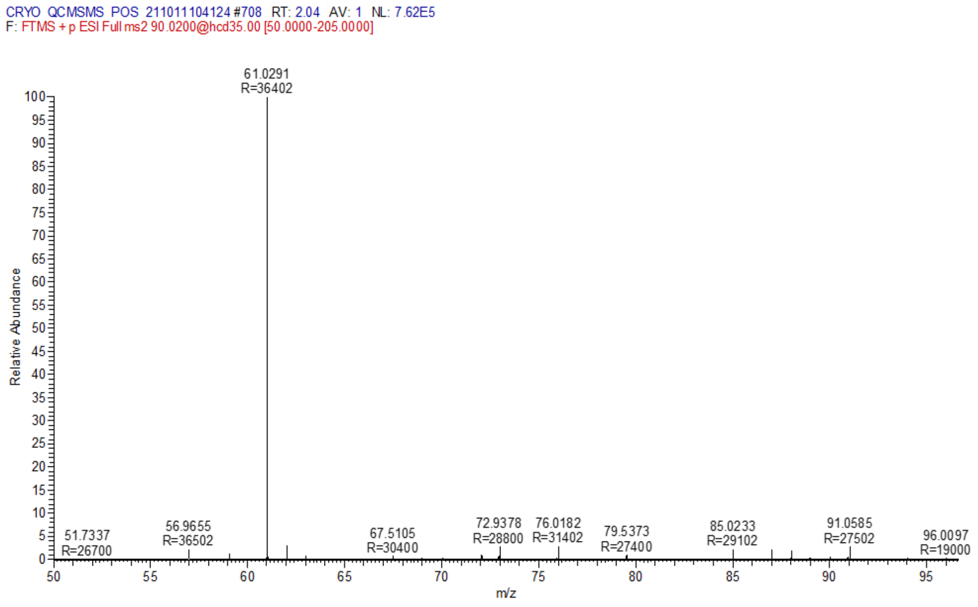


Experimental sample MS/MS spectra of m/z 90.0553 match the standard of L-Alanine.

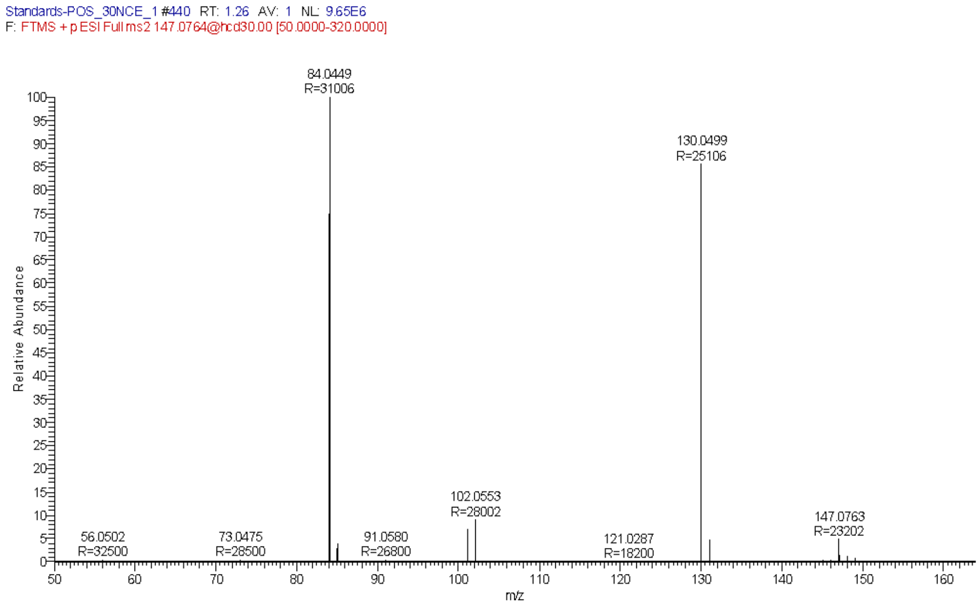


Experimental MS/MS spectra of the authentic standard of L-Glutamine.

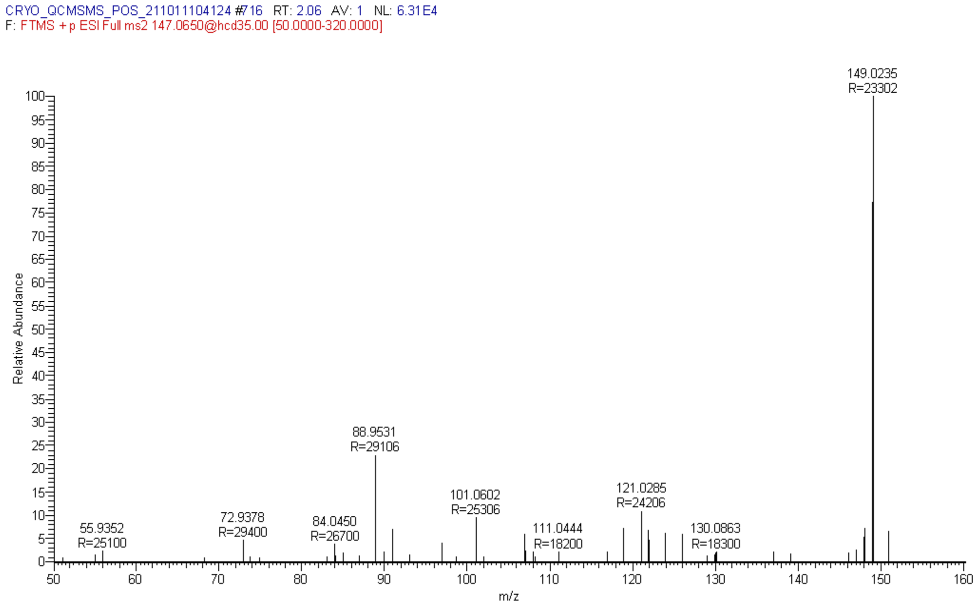


Experimental sample MS/MS spectra of m/z 147.0650 match the standard of L-Glutamine.

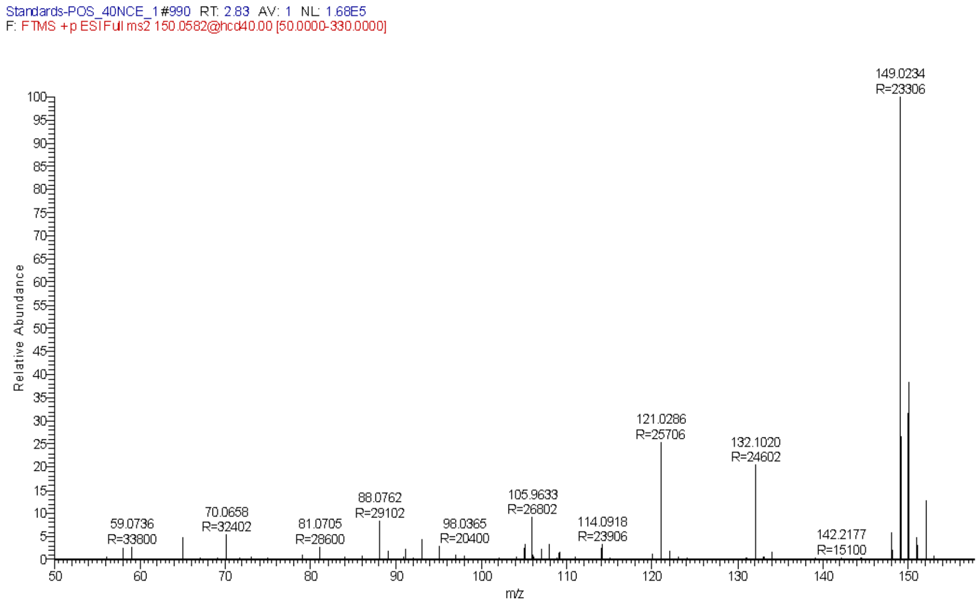


Experimental MS/MS spectra of the authentic standard of L-Methionine.

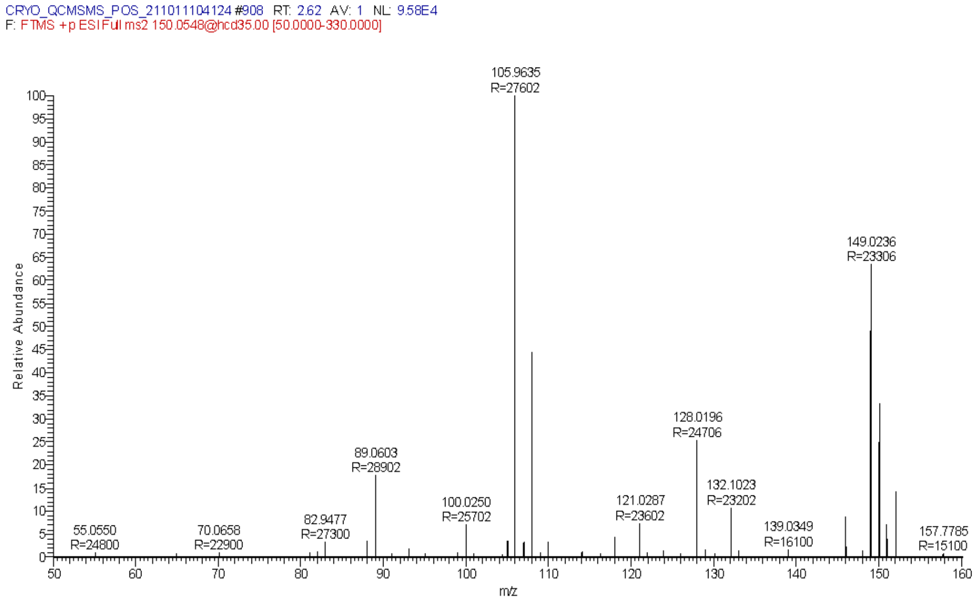


Experimental sample MS/MS spectra of m/z 150.0548 match the standard of L-Methionine.

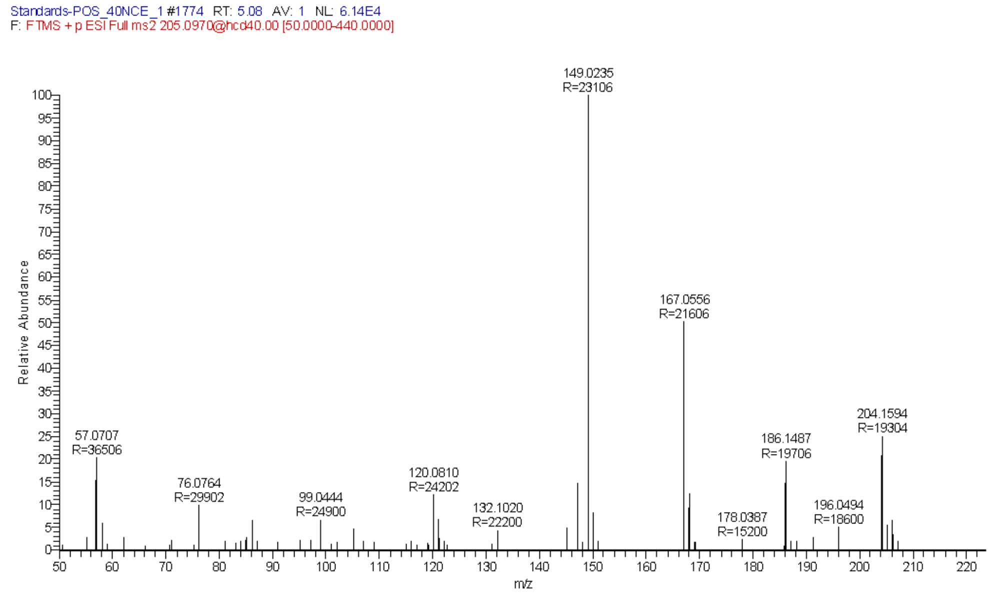


Experimental MS/MS spectra of the authentic standard of L-Tryptophan.

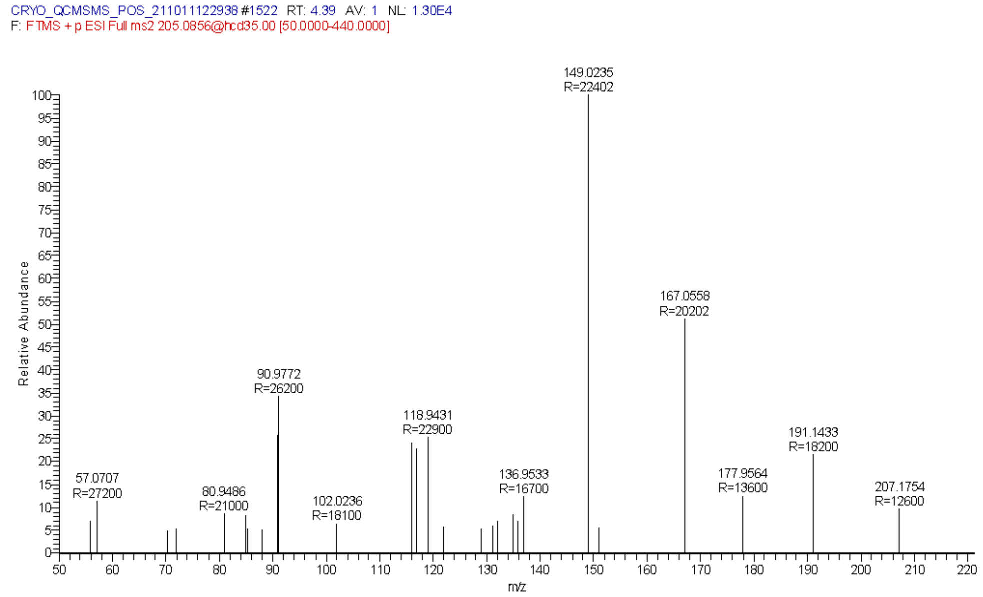


Experimental sample MS/MS spectra of m/z 205.0856 match the standard of L-Tryptophan.

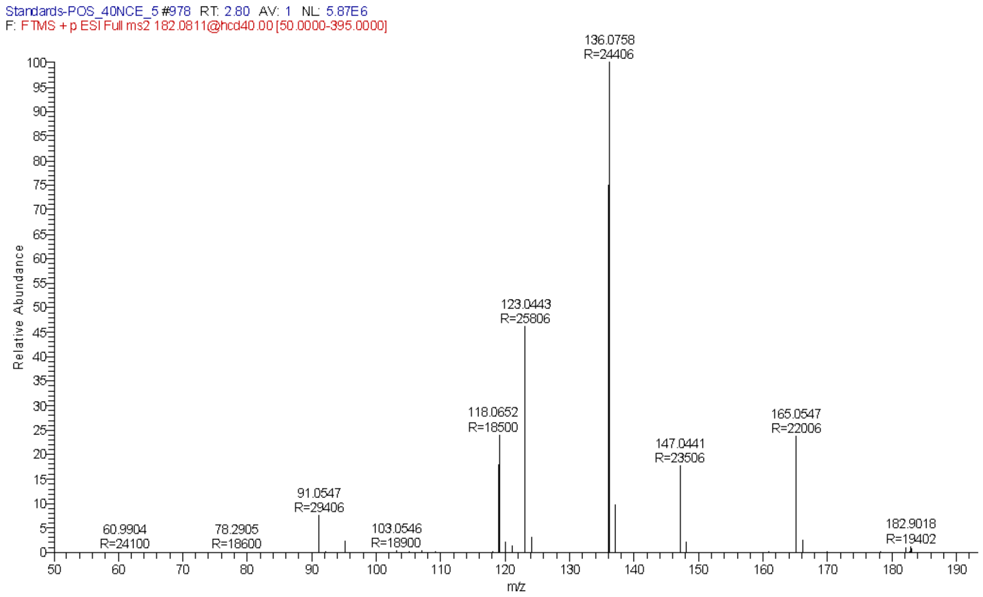


Experimental MS/MS spectra of the authentic standard of L-Tyrosine.

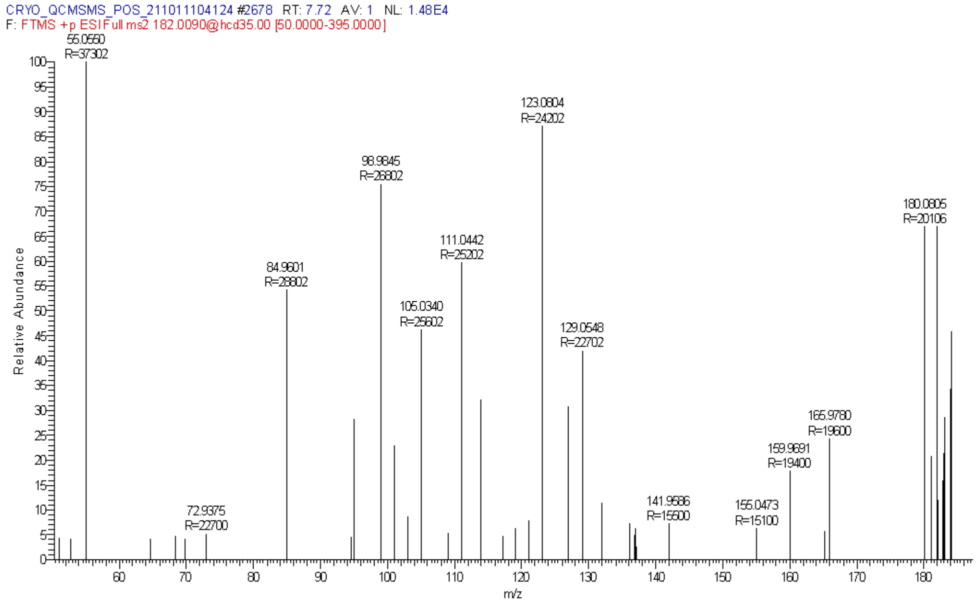


Experimental sample MS/MS spectra of m/z 182.0810 match with the standard of L-Tyrosine.

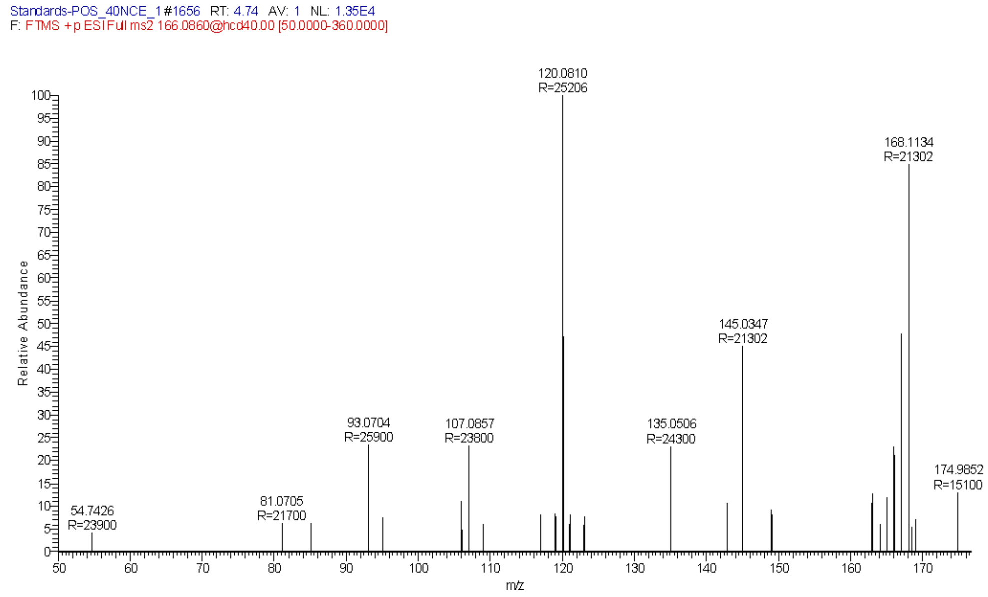


Experimental MS/MS spectra of the authentic standard of L-Phenylalanine.

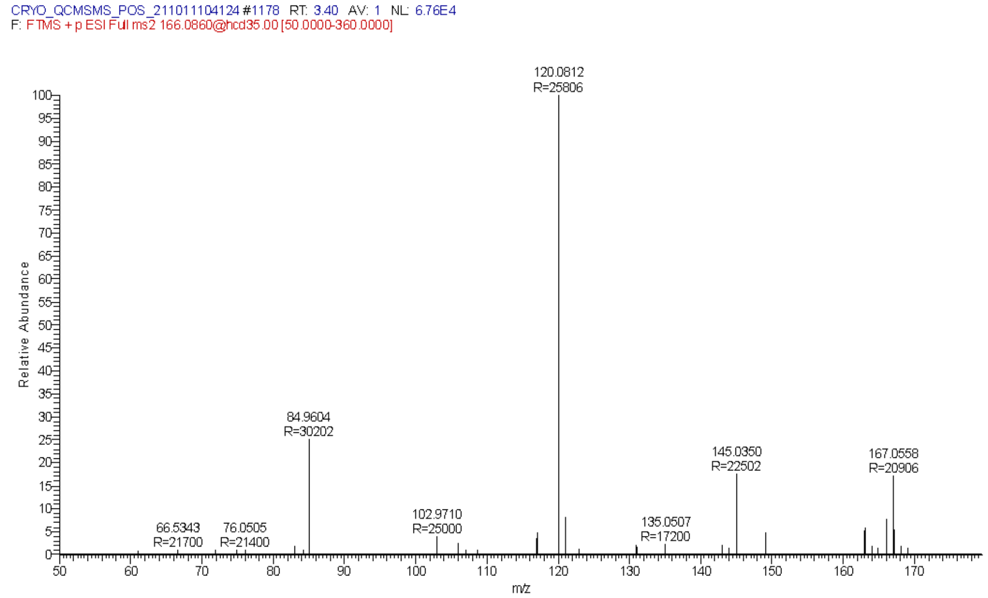


Experimental sample MS/MS spectra of m/z 166.0860 match the standard of L-Phenylalanine.

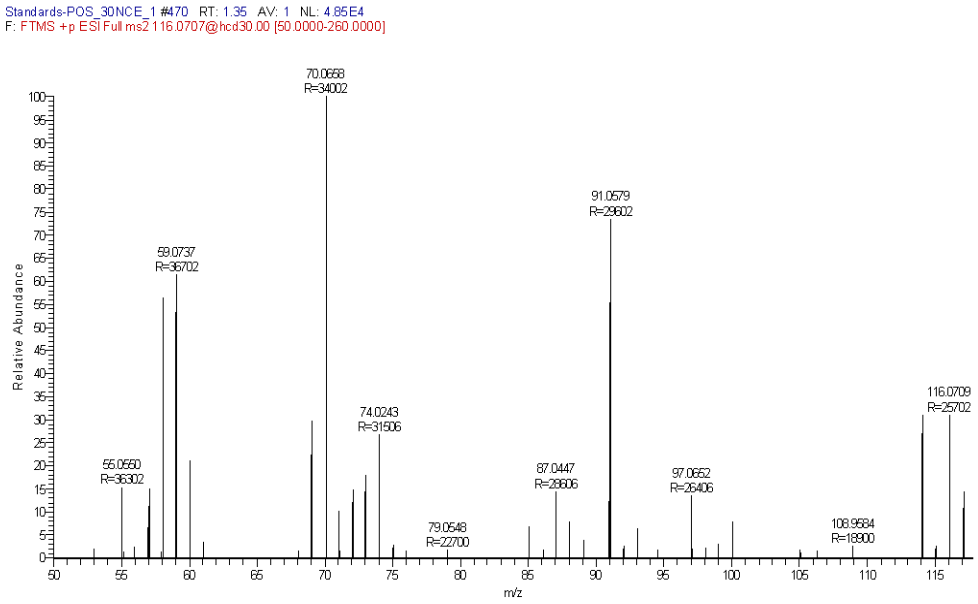


Experimental MS/MS spectra of the authentic standard of L-Proline.

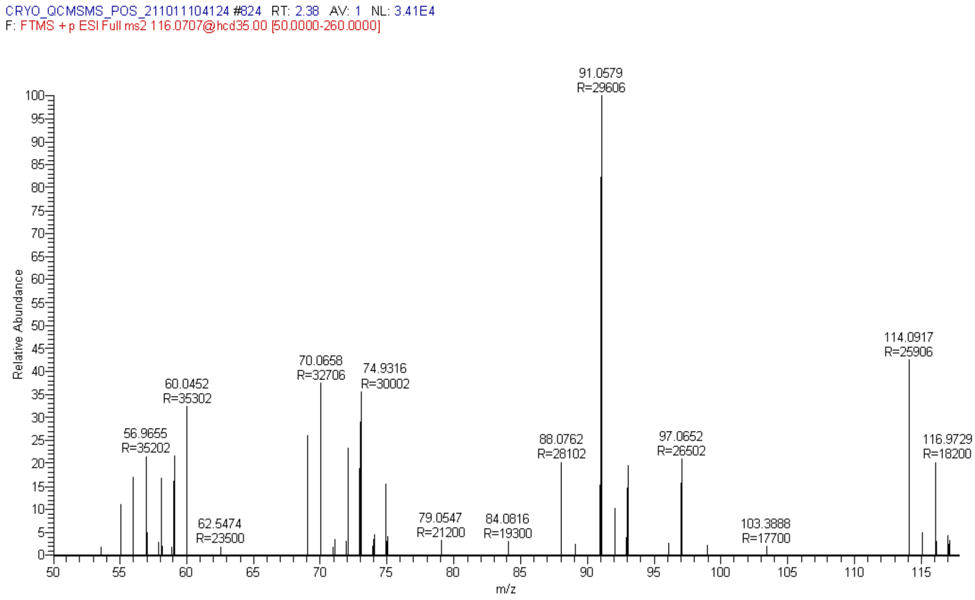


Experimental sample MS/MS spectra of m/z 116.0707 match the standard of L-Proline.

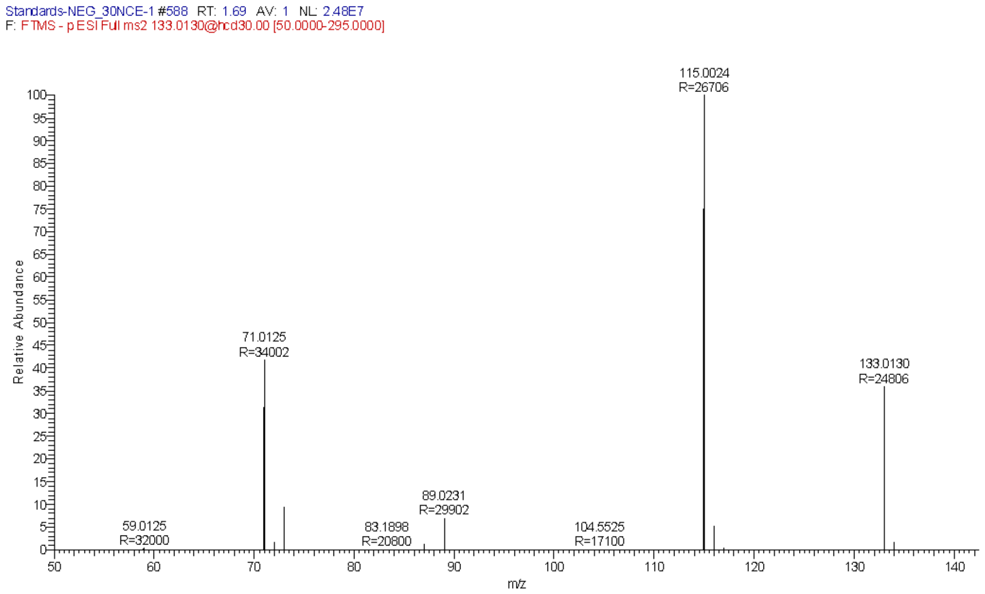


Experimental MS/MS spectra of the authentic standard of L-Malic acid.

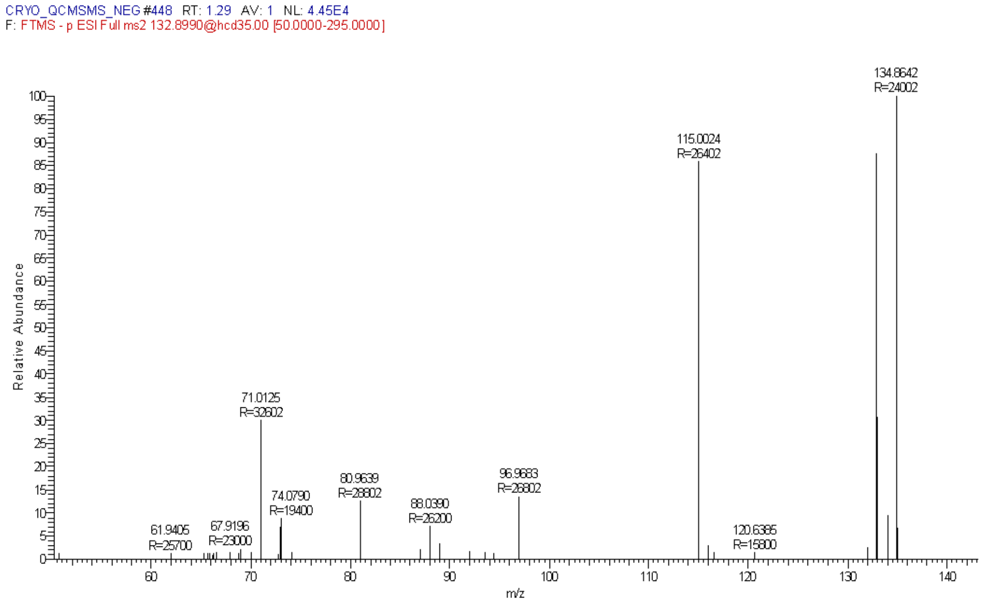


Experimental sample MS/MS spectra of m/z 133.0130 match the standard of L-Malic acid.

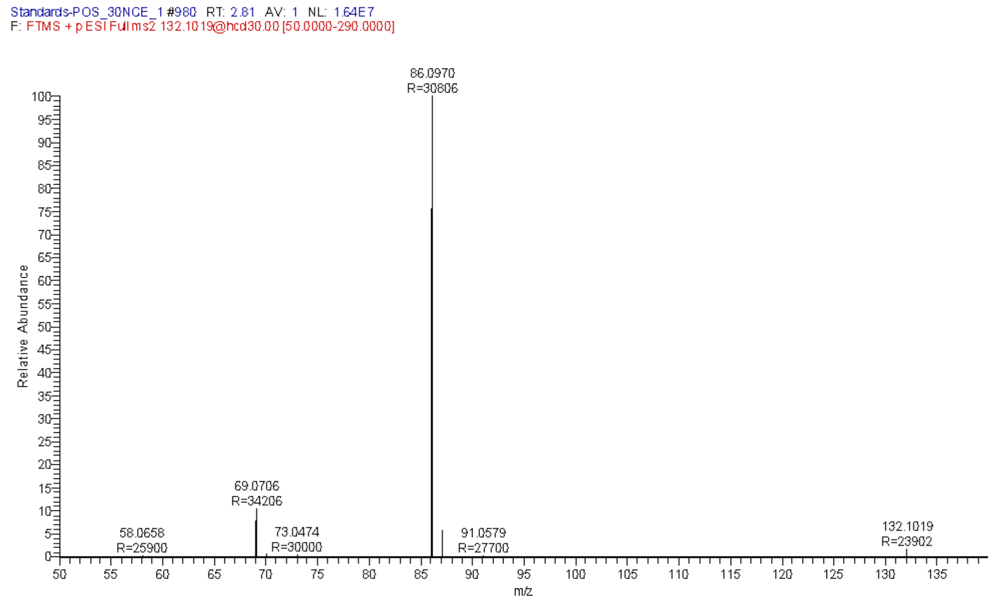


Experimental MS/MS spectra of the authentic standard of Isoleucine.

The

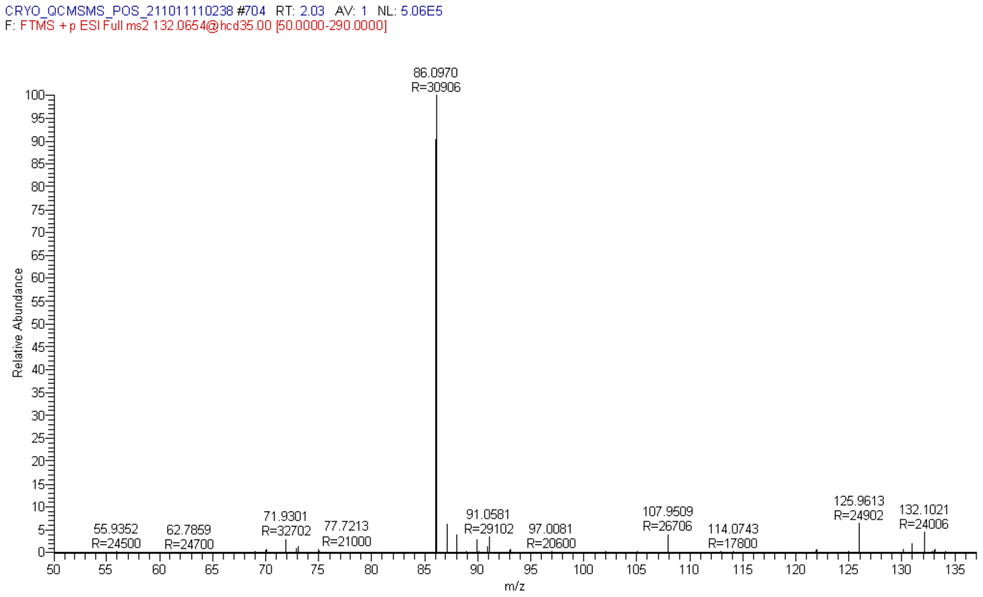


Experimental sample MS/MS spectra of m/z 132.0654 match the Isoleucine standard.

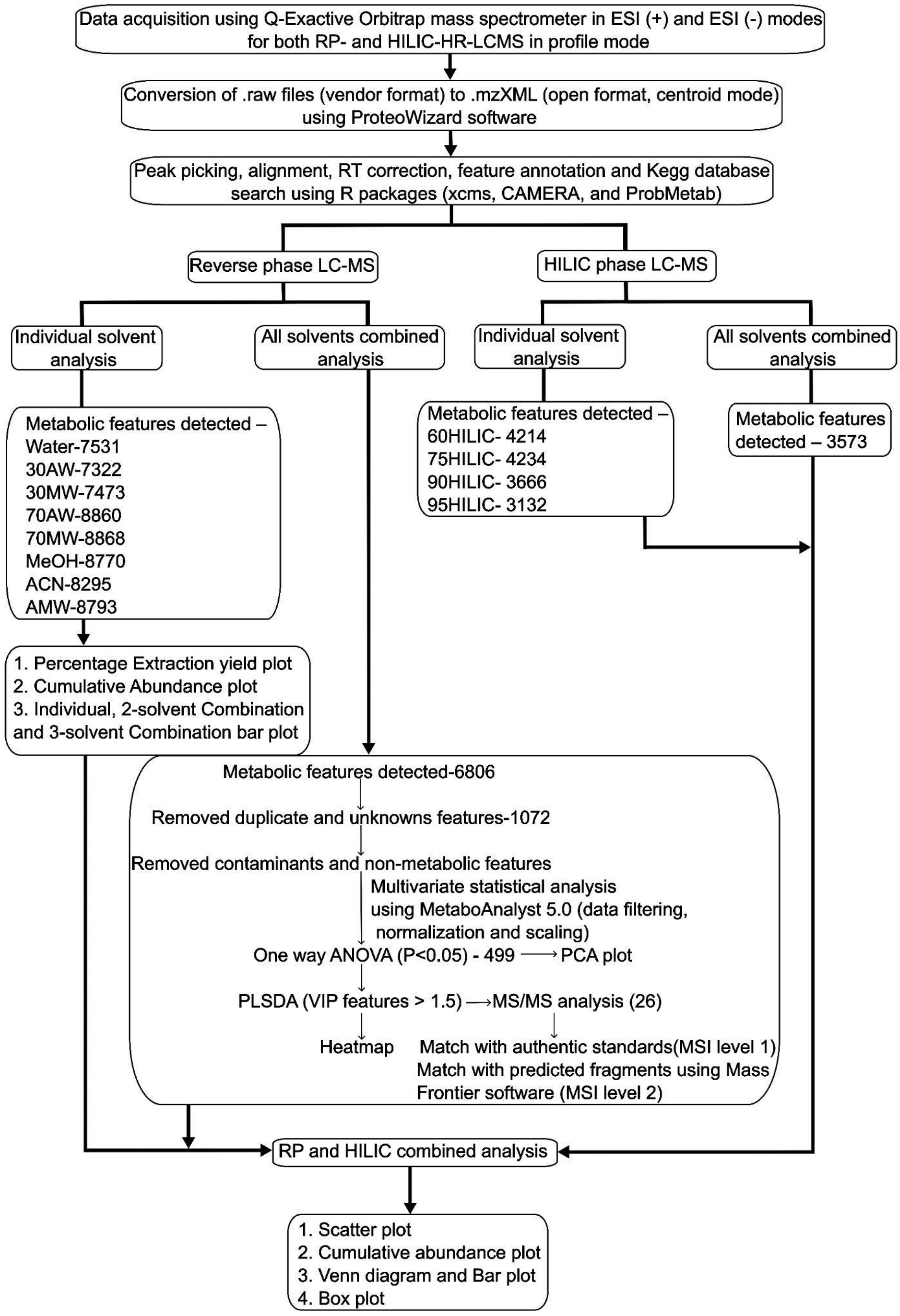


**Workflow S1: Workflow for data analysis for RP and HILIC data.**

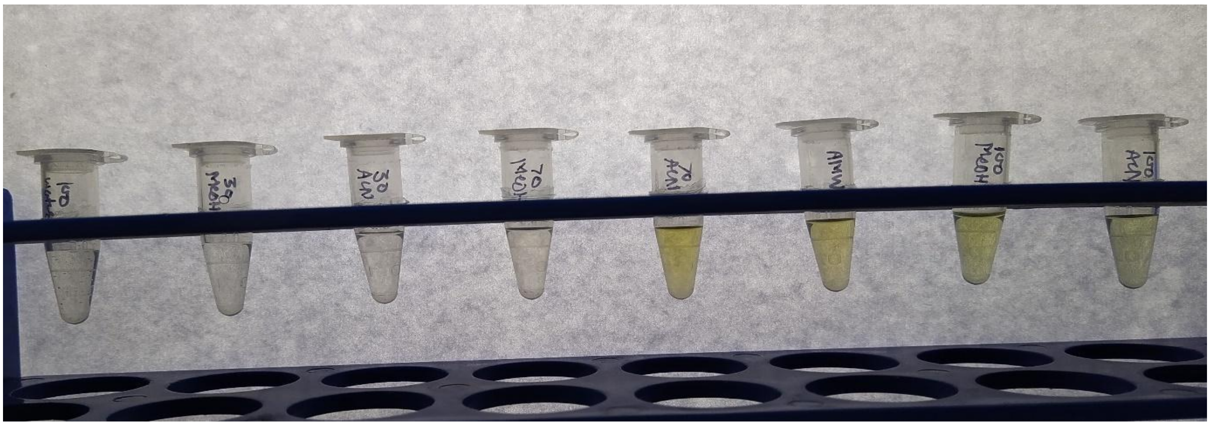

**Fig S5 (A): Visual inspection of extracts before drying.**

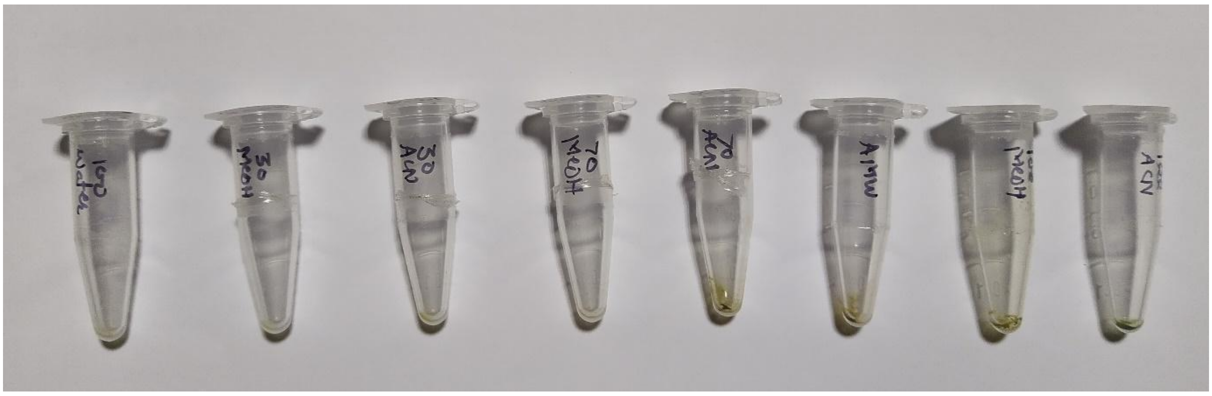

**Fig S5 (B): Visual inspection of extracts after drying.**

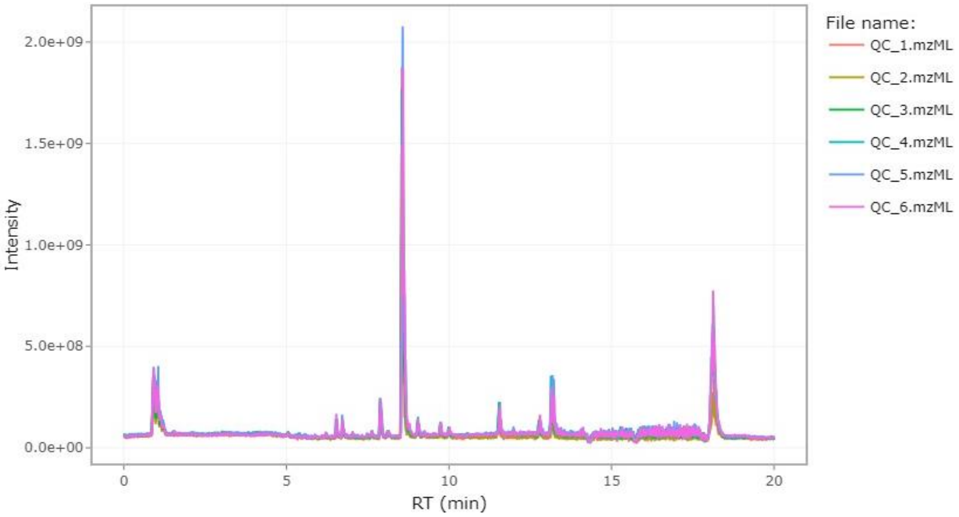


**Figure S6 (A): TIC Plot overlapping across QC samples.**

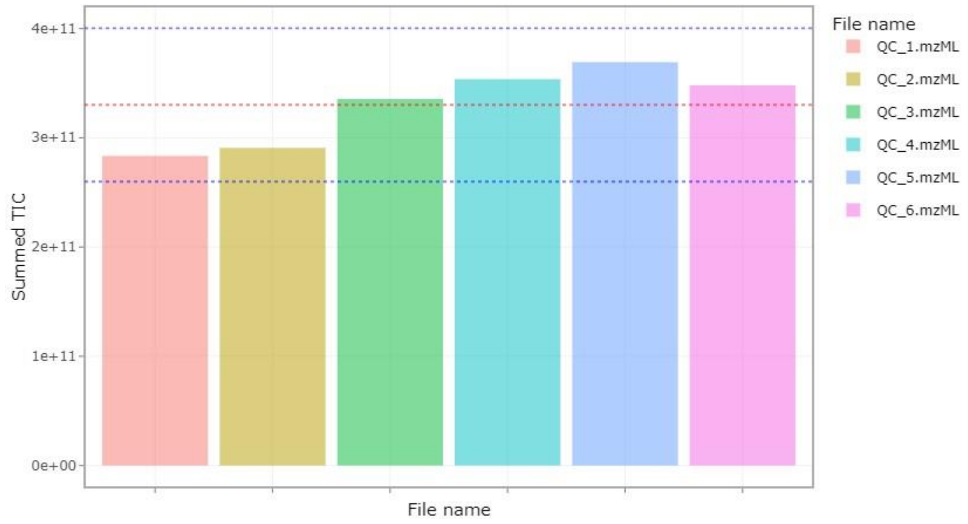


**Figure S6 (B): Summed TIC plot.**

The dashed red line is the mean of the summed TIC, and the blue lines represent the mean + 2SD and the mean - 2SD, respectively. Bars show the summed TIC of each QC file. The summed TIC variation between upper and lower dashed blue lines should be acceptable.

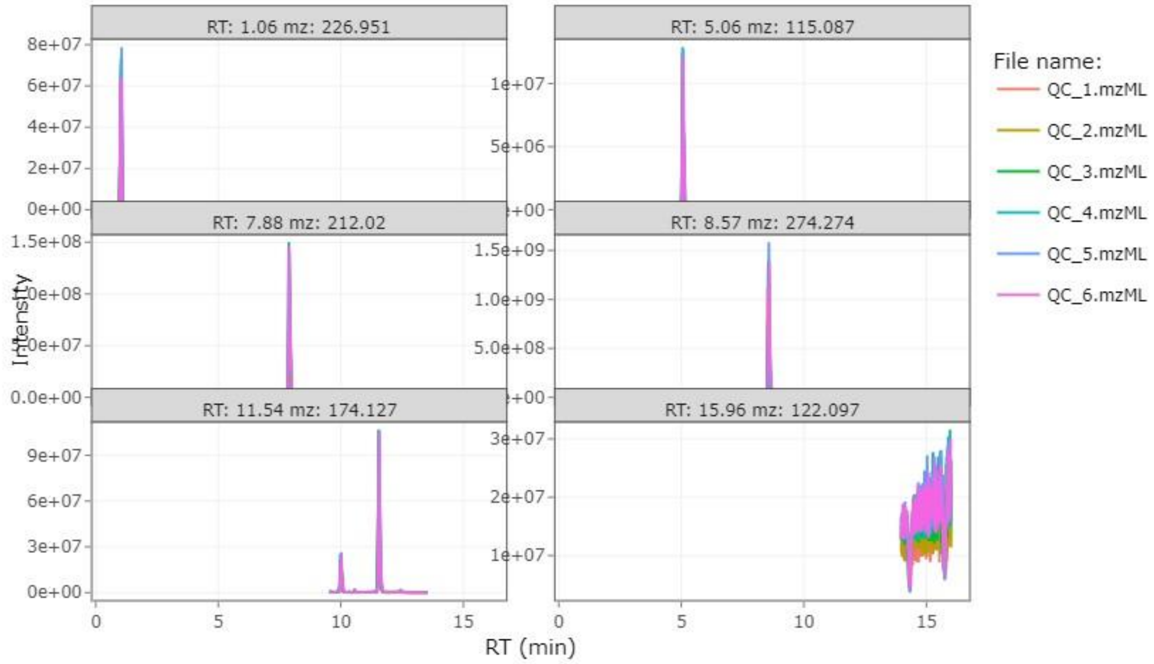


| Peak | Max RT Diff (min) | Max Mass Diff (ppm) | Max Intensity Ratio | Intensity CV (%) |
|---|---|---|---|---|
| RT: 1.06 mz: 226.951 | 0.04 | 0.94 | 1.47 | 13.6 |
| RT: 5.06 mz: 115.087 | 0.02 | 0.4 | 1.57 | 18.12 |
| RT: 7.88 mz: 212.02 | 0.04 | 0.65 | 1.45 | 14.12 |
| RT: 8.57 mz: 274.274 | 0.05 | 0.56 | 1.42 | 11.63 |
| RT: 11.54 mz: 174.127 | 0.05 | 0.26 | 1.55 | 19.35 |
| RT: 15.96 mz: 122.097 | 0.11 | 0.69 | 1.31 | 10.65 |

**Figure S7 (A): Summary of auto-selected peaks by RawHummus.**

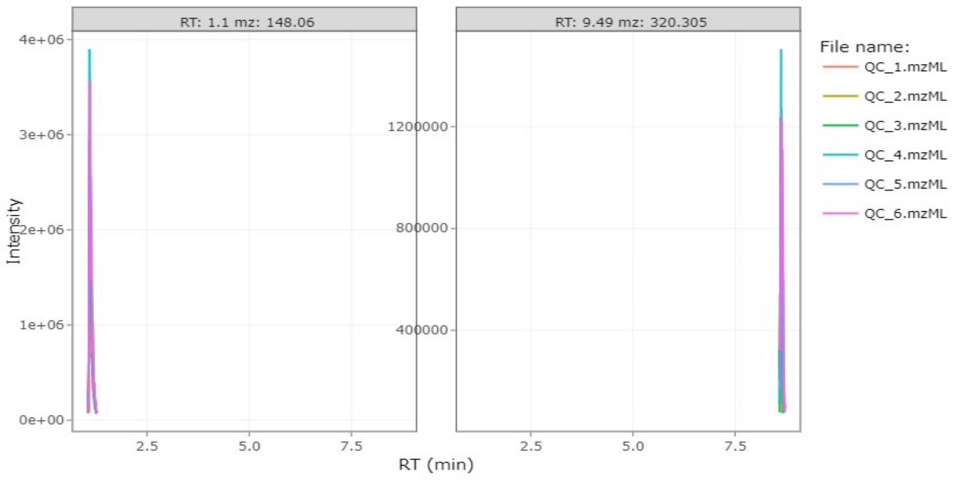


| Peak | Max RT Diff (min) | Max Mass Diff (ppm) | Max Intensity Ratio | Intensity CV (%) |
|---|---|---|---|---|
| RT: 1.1 mz: 148.06 | 0.01 | 0.52 | 1.58 | 16.73 |
| RT: 9.49 mz: 320.305 | 0.02 | 1.33 | 1.35 | 11.17 |

**Figure S7 (B): Summary of user-provided peaks.**

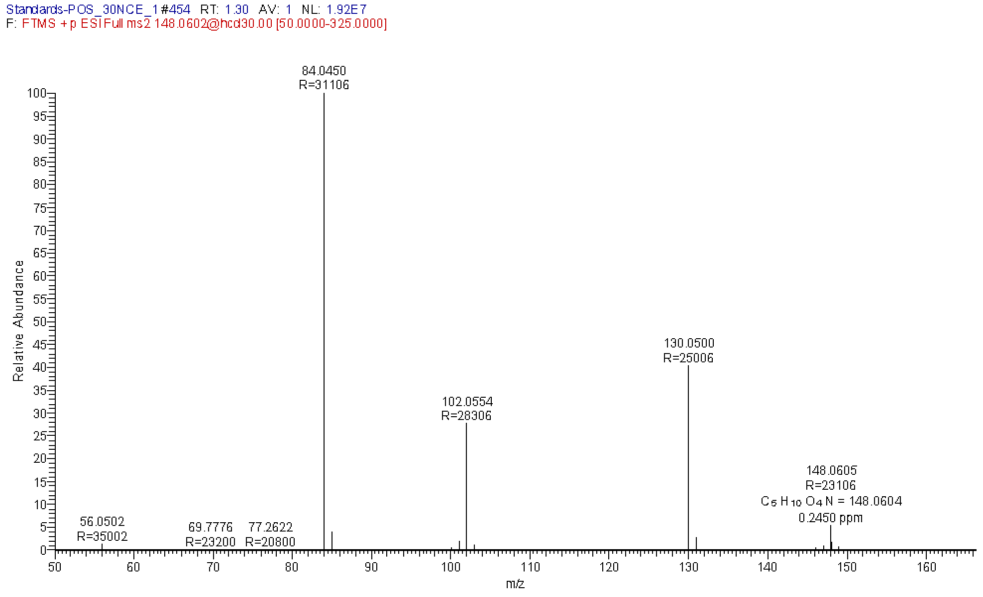


**Figure S7 (C): Experimental MS/MS spectra of the authentic standard of Glutamic acid.**

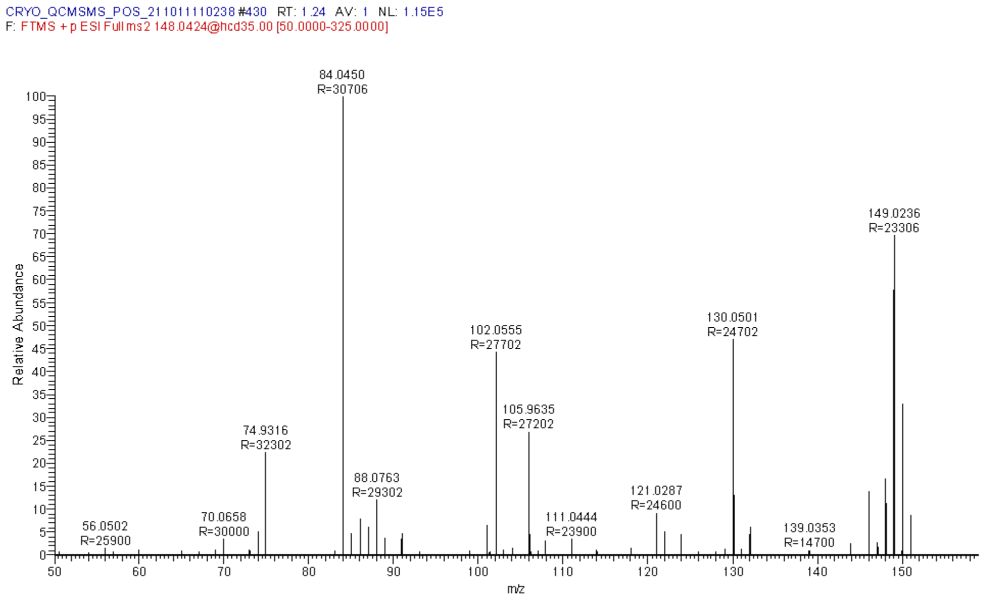


**Figure S7 (D): Experimental sample MS/MS spectra of m/z 148.0602 matching standard Glutamic acid MS/MS spectra.**

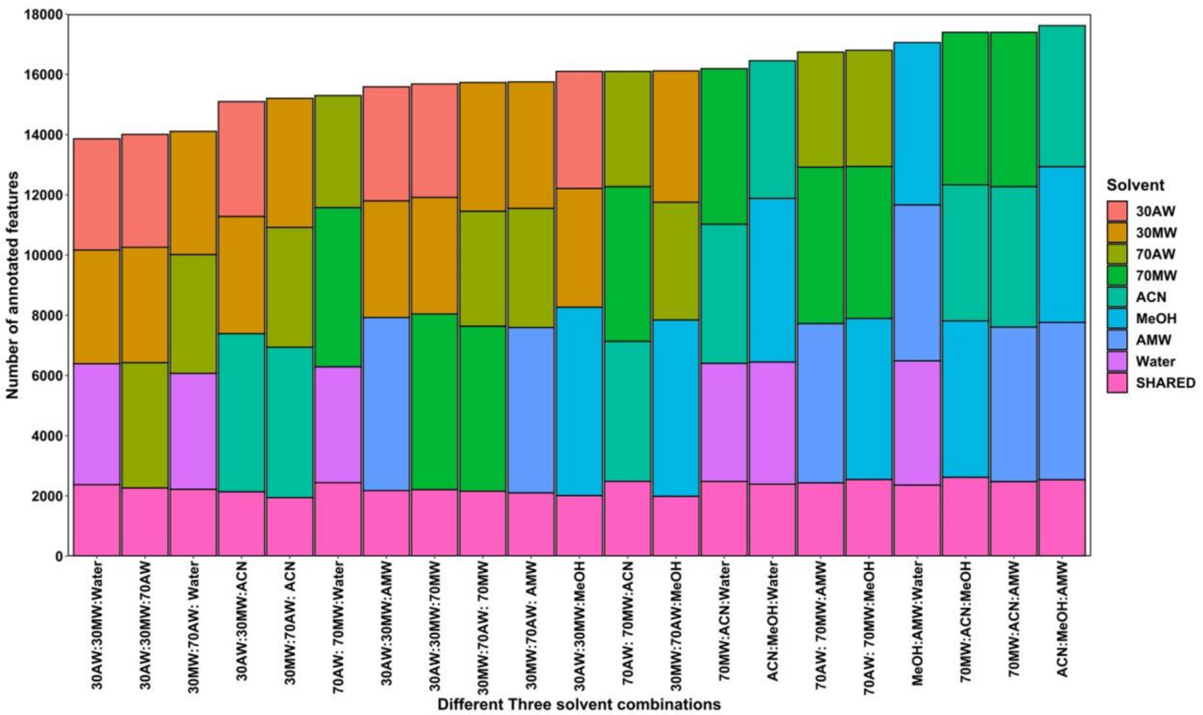


**Figure S8: Parallel 3-solvent combinations**

In three solvent combinations, solvents like ACN, MeOH, and AMW showed the tallest bar with the highest metabolic features. In a heatmap study, these solvents showed a similar pattern of metabolite extraction, contradicting 2-combination solvents, where complementary solvents were grouped. Solvents like 30AW, 30MW, and ACN showed minimum shared metabolic features and may show the highest chemical diversity. However, using three solvent systems in parallel combination or repeated extraction would make the study more laborious and cumbersome for large-scale studies.

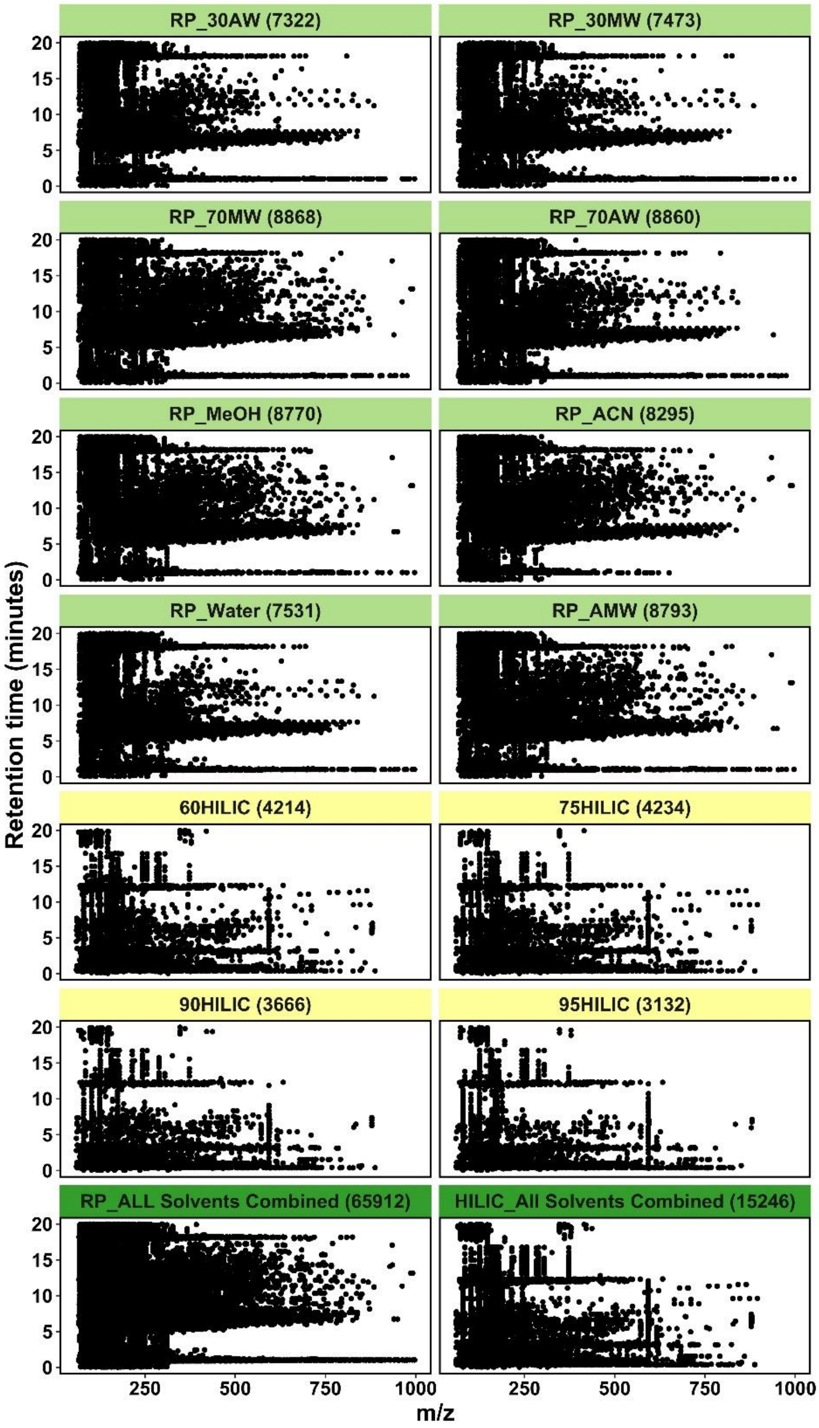


**Figure S9: RP and HILIC solvents scatter plot.**

Plot showing the distribution of metabolic features with retention time. Distribution is observed for different solvent extracts and RP and HILIC chromatography—individual features detected by black dots. A total number of ion features detected from individual solvents is given parenthetically.
