## Supplementary material for "Evaluation of extraction solvents for untargeted metabolomics to decipher the dissolved organic matter of Antarctic cryoconite holes": Supplimentary_Tables.pdf

### Table S1: Field-measured physicochemical characterization of cryoconite holes

| Sample ID | pH | Conductivity (µs) | ORP | TDS (mg/L) | Salinity (mg/L) | DO (mg/L) | Temperature (°C) |
|---|---|---|---|---|---|---|---|
| MC1 | 8.7 | 37.2 | -71 | 22.8 | 19.4 | 5.36 | 1.5 |
| MC2 | 8.62 | 13.4 | -92.3 | 8.1 | 6.4 | 5.54 | 1.3 |
| MC3 | 8.22 | 5.8 | -70.2 | 2.9 | 3.2 | 6.93 | 1.9 |
| MC4 | 7.38 | 14.2 | -45 | 9.5 | 7.3 | 6.11 | 2.3 |
| MC5 | 8.26 | 20.1 | -71.2 | 10.8 | 8.3 | 7.06 | 1.6 |
| MC6 | 8 | 15.5 | -51.3 | 10 | 7.8 | 6.8 | 1.3 |
| MC7 | 7.8 | 12.5 | -23.7 | 7.5 | 5.6 | 7.3 | 1.4 |
| MC8 | 7.74 | 15.5 | -17.2 | 10.2 | 7.8 | 7.17 | 1.3 |
| MC9 | 6.4 | 11.6 | 15.9 | 6.9 | 4.6 | 6.72 | 1.3 |
| MC10 | 6.91 | 17.8 | -6.5 | 12.2 | 9.3 | 7.06 | 1.3 |
| MC11 | 8.02 | 13.2 | -46.1 | 7.1 | 5.4 | 6.84 | 1.1 |
| MC12 | 7.9 | 11.6 | 11.6 | 7.8 | 5.8 | 7.12 | 1.4 |
| MC13 | 7.64 | 19 | -44.5 | 12 | 9.2 | 6.56 | 1.9 |
| MC14 | 7.7 | 12 | -41.5 | 7.3 | 5 | 7.13 | 1.8 |
| MC15 | 7.8 | 15 | -54 | 9.3 | 7 | 7.17 | 2.3 |
| MC16 | 6.68 | 45.1 | -17.8 | 28.8 | 21.9 | 7.89 | 1.9 |
| MC17 | 7.18 | 28 | -5.4 | 16.6 | 12.3 | 7.35 | 1.3 |
| MC18 | 7.15 | 35.4 | -5.7 | 22 | 17 | 7 | 1.7 |
| MC19 | 7.14 | 43.2 | -23.8 | 27.1 | 20.5 | 6.86 | 1.4 |
| MC20 | 7.29 | 45.1 | -17.8 | 29.2 | 21.8 | 7.13 | 1.6 |

### Table S7: List of authentic standards (low molecular weight organic compounds) used to evaluate untargeted, high-resolution mass spectrometry technique

| Compound | Class | Kegg ID | Formula | Monoisotopic mass | Observed ion $[M+H]^+$ | Mass accuracy (ppm) |
|---|---|---|---|---|---|---|
| L-Tyrosine | Amino acid | C00082 | C9H11NO3 | 181.0739 | 182.0810 | -0.72 |
| L-Isoleucine | Amino acid | C00407 | C6H13NO2 | 131.0946 | 132.1019 | -017 |
| L-Histidine | Amino acid | C00135 | C6H9N3O2 | 155.0695 | 156.0767 | -0.60 |
| L-Tryptophan | Amino acid | C00078 | C11H12N2O2 | 204.0899 | 205.0969 | -1.05 |
| L-Asparagine | Amino acid | C00152 | C4H8N2O3 | 132.0535 | 133.0604 | -2.8 |
| L-Methionine | Amino acid | C00073 | C5H11NO2S | 149.051 | 150.0582 | -0.66 |
| L-Glutamic acid | Amino acid | C00025 | C5H9NO4 | 147.0532 | 148.0603 | -0.99 |
| L-Cysteine | Amino acid | C00097 | C3H7NO2S | 121.0197 | 122.0272 | 1.04 |
| L-Threonine | Amino acid | C00188 | C4H9NO3 | 119.0582 | 120.0657 | 1.09 |
| L-Phenylalanine | Amino acid | C00079 | C9H11NO2 | 165.079 | 166.0861 | -0.72 |
| L-Proline | Amino acid | C00148 | C5H9NO2 | 115.0633 | 116.0707 | 1.16 |

| | | | | | | |
|---|---|---|---|---|---|---|
| L-Glutamine | Amino acid | C00064 | C5H10N2O3 | 146.0691 | 147.0763 | -0.64 |
| L-Serine | Amino acid | C00065 | C3H7NO3 | 105.0426 | 106.0502 | 2.7 |
| L-Alanine | Amino acid | C00041 | C3H7NO2 | 89.0477 | 90.0554 | 4.6 |
| L-Valine | Amino acid | C00183 | C5H11NO2 | 117.079 | 118.0864 | 1.5 |
| L-Leucine | Amino acid | C00123 | C6H13NO2 | 131.0946 | 132.1019 | -017 |
| L-Glutathione | Peptide | C00051 | C10H17N3O6 S | 307.0838 | 308.0907 | -1.34 |
| D-Mannitol | Carbohydrate | C00392 | C6H14O6 | 182.079 | 183.0862 | -0.8 |
| D-Sucrose | Carbohydrate | C00089 | C12H22O11 | 342.1162 | 343.1229 | -1.6 |
| Pyridoxine Hydrochloride | Vitamin | C00314 | C8H11NO3 | 169.0739 | 170.0810 | -0.9 |
| Riboflavin | Vitamin | C00255 | C17H20N4O6 | 376.1383 | 377.1449 | -1.68 |
| L-Malic acid** | Organic acid | C00149 | C4H6O5 | 134.0215 | 133.0130 | -0.78 |

** L-Malic acid is detected in negative ESI mode $[M-H]^-$

**Table S8: Metabolic features putatively confirmed by MS/MS spectral fragments match with the theoretical fragments from Mass Frontier (7.0, Thermo Fisher) software**

| KEGG ID | Compound Name | Formula | Adduct | Mono isotopic mass | Observed m/z | RT (min) | ppm error | MS/MS fragments match |
|---|---|---|---|---|---|---|---|---|
| C00250 | Pyridoxal | C8H9NO3 | M+H | 167.0582 | 168.0653 | 5.7 | 1 | 123.04 |
| C00262 | Hypoxanthine | C5H4N4O | M+H | 136.0385 | 137.0456 | 2.6 | 1 | 67.03,82.04,121.05 |
| C00319 | Sphingosine | C18H37NO2 | M+H | 299.2824 | 300.2891 | 8.8 | 2 | 69.03,107.09,129.09,252.27 |
| C00334 | 4-Aminobutanoate | C4H9NO2 | M+H | 103.0633 | 104.0708 | 1.0 | 2 | 58.07, 70.07 |
| C00387 | Guanosine | C10H13N5O5 | M+H | 283.0917 | 284.0984 | 2.2 | 2 | 85.03, 203.06 |
| C00437 | N-Acetylornithine | C7H14N2O3 | M+H | 174.1004 | 175.1075 | 1.1 | 1 | 57.03, 84.04, 143.08 |
| C00475 | Cytidine | C9H13N3O5 | M+H | 243.0855 | 244.0923 | 1.1 | 2 | 85.03,129.07,194.06 |
| C00642 | 3-Hydroxyphenylacetate | C8H8O3 | M+H | 152.0473 | 153.0544 | 6.3 | 1 | 107.05, 137.06 |
| C00864 | Pantothenic acid | C9H17NO5 | M+H | 219.1107 | 220.1175 | 4.8 | 2 | 101.06,112.04,129.05 |
| C01879 | 5-oxoproline | C5H7NO3 | M+H | 129.0426 | 130.0498 | 1.1 | 1 | 71.01,102.05 |
| C02727 | N6-Acetyl-L-lysine | C8H16N2O3 | M+H | 188.1161 | 189.1231 | 1.1 | 1 | 88.04,125.11 |
| C06180 | Anabasine | C10H14N2 | M+H | 162.1157 | 163.1227 | 1.6 | 2 | 80.05,84.08,106.07,118.07 |
| C06231 | Ectoine | C6H10N2O2 | M+H | 142.0742 | 143.0813 | 1.1 | 1 | 72.01,102.05,125.07 |
| C06577 | Cuminaldehyde | C10H12O | M+H | 148.0888 | 149.0957 | 13.1 | 3 | 79.05,107.05 |
| C10048 | Ginkgetin | C32H22O10 | M+H | 566.1213 | 567.1307 | 11.3 | 4 | 167.03,269.04 |

**Table S9: TIC correlation analysis and metabolic profile similarity:**

| QC Samples | QC_1. mzML | QC_2. mzML | QC_3. mzML | QC_4. mzML | QC_5. mzML | QC_6. mzML |
|---|---|---|---|---|---|---|
| QC_1. mzML | 1 | 0.996 | 0.944 | 0.959 | 0.938 | 0.914 |
| QC_2. mzML | 0.996 | 1 | 0.957 | 0.969 | 0.949 | 0.929 |
| QC_3. mzML | 0.944 | 0.957 | 1 | 0.996 | 0.99 | 0.983 |
| QC_4. mzML | 0.959 | 0.969 | 0.996 | 1 | 0.989 | 0.983 |
| QC_5. mzML | 0.938 | 0.949 | 0.99 | 0.989 | 1 | 0.996 |
| QC_6. mzML | 0.914 | 0.929 | 0.983 | 0.983 | 0.996 | 1 |

**Table S10: Extraction solvents for different metabolomics studies**

| Reference | Analytical Platform | Extraction solvents and their combinations (v/v) evaluated | Best solvent and combinations (v/v) suggested/recommended | Matrix/Sample for extraction |
|---|---|---|---|---|
| Villas-Boas et al., 2005 | GC-MS | $CHCl_3$: MeOH: Buffer (CMB), boiling EtOH, perchloric acid, potassium hydroxide, MeOH: water, MeOH | MeOH | Saccharomyces cerevisiae culture |
| Winder C et al., 2008 | GC-MS | Perchloric acid, 100% MeOH, MeOH: $CHCl_3$ (2:1), 0.25 M KOH, boiling EtOH | MeOH | E. coli culture |
| Bruce et al., 2009 | LC-MS | Acetone, MeOH, ACN, and EtOH and their combinations | MeOH: EtOH (1:1) and MeOH: ACN: acetone (1:1:1) | Human Blood plasma |
| Theodoridis et al., 2011 | LC-MS | MeOH: water: $CHCl_3$ (different combinations) | MeOH: $CHCl_3$: water (40:40:20) | Vitis vinifera (grapes) |
| Kim et al., 2013 | GC-MS | 100% MeOH, ACN: water (1:1), ACN: MeOH: Water (2:2:1), boiling EtOH (75%) | ACN: water (1:1) at −20 °C | Saccharomyces cerevisiae culture |
| Martin et al., 2014 | LC-MS | Hexane, DCM, ethyl acetate, MeOH, IPA, water, EtOH: water (70:30); Solvent partitioning | 70% EtOH | Aerial tissue (dried berries, leaves, and stem) from three different plant species |
| Swenson et al., 2015 and 2018 | GC- and LC-MS | Water, 10-100% MeOH in water, 10mM $K_2SO_4$, 10mM $NH_4HCO_3$, IPA: MeOH: water (3:3:2), ethyl acetate | Water for polar and Ethyl acetate for non-polar metabolites | Rhizopheric soil |
| Doppler et al., 2016 | LC-MS | Aqueous mixtures of MeOH, ACN, MeOH: ACN: Water | MeOH: ACN: water (3.75:3.75:2.5) | Wheat organs |

| | | | | |
|---|---|---|---|---|
| Deda et al., 2016 | NMR, LC- and GC-MS | Aqueous mixtures of MeOH, IPA, and ACN | Aqueous ACN (1:1) | Rat feces |
| Moosmang et al., 2017 | NMR and LC-MS | Water, MeOH, ACN, MeOH: water (4:1), ACN: water (1:3), ethyl acetate, $CHCl_3$: MeOH: water (3:1:2) | Water | Human feces |
| Cheng et al., 2020 | LC-MS | MeOH, ACN, and water in different ratios | MeOH | Human feces |
| Schippers et al., 2023 | LC-MS | 100% MeOH, ACN: MeOH: water (5:3:2), MeOH: water: $CHCl_3$ (2:1:2 and 1:2:2) | MeOH | Zebrafish larvae |
| Srivastava et al., 2023 | GC-MS | MeOH, ACN, MeOH: ACN (50:50), MeOH: MeOH: water (MMW), $CHCl_3$ | MeOH: MeOH: water (MMW) | Human Plasma |
| Hemmer et al., 2024 | LC-MS | MeOH, ACN, water, and their combinations | MeOH | Human and Rat urine |
| Navarrete-Carriola et al., 2024 | LC- and GC-MS | Water, MeOH, hexane, and DCM | MeOH | Roots and Leaves of *Iostephane heterophylla* |

Abbreviations: MeOH (Methanol), EtOH (Ethanol), ACN(Acetonitrile), IPA(Isopropanol), DCM(Dichloromethane), KOH (Potassium hydroxide), $K_2SO_4$(Potassium sulfate, $NH_4HCO_3$ (Ammonium bicarbonate), $CHCl_3$ (Chloroform)

**Tabel S11: Solvents and their strength and polarity indices (adapted from Snyder 1978)**

| Solvent | Selectivity group | Solvent strength (solubility power) | P′ | Xe | Xd | Xn |
|---|---|---|---|---|---|---|
| Acetonitrile | VIa (nitriles) | 0.65 | 6.2 | 0.3 | 0.26 | 0.41 |
| Methanol | II (aliphatic alcohols) | 0.95 | 6.6 | 0.51 | 0.19 | 0.3 |
| Water | VIII (water) | >>1 | 9 | 0.4 | 0.34 | 0.26 |

P′ is the polarity index of the solvent, and Xe, Xd, and Xn represent the fraction of P′ contributed by interactions associated with proton-donor, proton-acceptor, and dipole moment characteristics of the solvent, respectively.

**Table S12: Biologically important metabolites (putatively annotated) that are detected in RP and HILIC or both techniques in positive or negative ESI mode**

| KEGG ID | Metabolite Name | Proposed Mass | Ion annotation | ESI mode | Technique | Class | Subclass_1 | Subclass_2 | Subclass_3 |
|---|---|---|---|---|---|---|---|---|---|
| C02727 | N6-Acetyl-L-lysine | 188.1157 | 187.1082 | Negative | HILIC | Acylated amino acid | NA | NA | NA |

| | | | | | | | | | |
|---|---|---|---|---|---|---|---|---|---|
| C00116 | Glycerol | 92.0472 | 93.0550 | Positive | Both | Carbohydrates | Monosaccharides | Sugar alcohols | NA |
| C00031 | D-Glucose | 180.0627 | 179.0554 | Negative | HILIC | Carbohydrates | Monosaccharides | Aldoses | NA |
| C00089 | Sucrose | 342.1165 | 341.1092 | Negative | HILIC | Carbohydrates | Oligosaccharides | Disaccharides | NA |
| C00794 | D-Sorbitol | 182.0795 | 217.0481 | Negative | HILIC | Carbohydrates | Monosaccharides | Sugar alcohols | NA |
| C01083 | Trehalose | 342.1150 | 343.1228 | Positive | RP | Carbohydrates | Oligosaccharides | Disaccharides | NA |
| C08261 | Azelaic acid | 188.1043 | 187.0970 | Negative | Both | Lipids | Fatty acyls | Fatty Acids and Conjugates | Dicarboxylic acids |
| C08491 | (-)-Jasmonic acid | 210.1253 | 211.1325 | Positive | RP | Lipids | Fatty acyls | Octadecanoids | jasmonic acids |
| C08490 | Jasmone | 164.1193 | 165.1272 | Positive | RP | Lipids | Fatty acyls | Octadecanoids | jasmonic acids |
| C11512 | Methyl jasmonate | 224.1403 | 225.1481 | Positive | RP | Lipids | Fatty acyls | Octadecanoids | jasmonic acids |
| C08261 | Azelaic acid | 188.1056 | 189.1117 | Positive | Both | Lipids | Fatty acyls | Fatty Acids and Conjugates | Dicarboxylic acids |
| C08491 | (-)-Jasmonic acid | 210.1253 | 211.1325 | Positive | RP | Lipids | Fatty acyls | Octadecanoids | jasmonic acids |
| C08490 | Jasmone | 164.1193 | 165.1272 | Positive | RP | Lipids | Fatty acyls | Octadecanoids | jasmonic acids |
| C11512 | Methyl jasmonate | 224.1403 | 225.1481 | Positive | RP | Lipids | Fatty acyls | Octadecanoids | jasmonic acids |
| C06103 | 6-Hydroxyhexanoic acid | 132.0779 | 133.0858 | Positive | RP | Lipids | Fatty acyls | Fatty Acids and Conjugates | Hydroxy fatty acids |
| C14828 | 9,10-DHOME | 314.2445 | 315.2524 | Positive | RP | Lipids | Fatty acyls | Fatty Acids and Conjugates | Hydroxy fatty acids |
| C14774 | 11,12-DHET | 338.2444 | 339.2523 | Positive | RP | Lipids | Fatty acyls | Eicosanoids | Hydroxy/hydroperoxyeicosatrienoic acids |
| C14826 | 12(13)-EpOME | 296.2341 | 297.2419 | Positive | RP | Lipids | Fatty acyls | Octadecanoids | Other Octadecanoids |
| C14762 | 13(S)-HODE | 296.2340 | 297.2418 | Positive | RP | Lipids | Fatty acyls | Octadecanoids | Other Octadecanoids |
| C14825 | 9(10)-EpOME | 296.2339 | 297.2417 | Positive | RP | Lipids | Fatty acyls | Octadecanoids | Other Octadecanoids |
| C14827 | 9(S)-HPODE | 312.2269 | 313.2348 | Positive | RP | Lipids | Fatty acyls | Octadecanoids | Other Octadecanoids |
| C16322 | 9-Oxononanoic acid | 172.1125 | 158.0962 | Positive | RP | Lipids | Fatty acyls | Fatty Acids and Conjugates | Oxo fatty acids |
| C01092 | 8-Amino-7-oxononanoate | 187.1188 | 188.1293 | Positive | RP | Lipids | Fatty acyls | Fatty Acids and Conjugates | Oxo fatty acids |
| C06761 | (4E)-2-Oxohexenoic acid | 128.0476 | 151.0350 | Positive | RP | Lipids | Fatty acyls | Fatty Acids and Conjugates | Oxo fatty acids |
| C00164 | Acetoacetate | 102.0314 | 103.0392 | Positive | RP | Lipids | Fatty acyls | Fatty Acids and Conjugates | Oxo fatty acids |
| C17224 | 2-Oxo-8-methylthiooctanoic acid | 204.0839 | 205.0917 | Positive | RP | Lipids | Fatty acyls | Fatty Acids and Conjugates | Oxo fatty acids |
| C12027 | 2-Amino-9,10-epoxy-8-oxodecanoic acid | 215.1150 | 216.1228 | Positive | RP | Lipids | Fatty acyls | Fatty Acids and Conjugates | Oxo fatty acids |
| C13846 | Octadecanamide | 283.2869 | 284.2941 | Positive | RP | Lipids | Fatty acyls | Fatty amides | Primary amides |
| C13831 | Dodecanamide | 199.1928 | 200.2007 | Positive | RP | Lipids | Fatty acyls | Fatty amides | Primary amides |
| C00319 | Sphingosine | 299.2818 | 300.2891 | Positive | RP | Lipids | Sphingolipids | Sphingoid bases | Sphing-4-enines (Sphingosines) |
| C00836 | Sphinganine | 301.2975 | 302.3048 | Positive | RP | Lipids | Sphingolipids | Sphingoid bases | Sphinganines |
| C06124 | Sphingosine 1-phosphate | 379.2432 | 380.2510 | Positive | RP | Lipids | Sphingolipids | Sphingoid bases | Sphingoid base 1-phosphates |
| C01747 | Psychosine | 461.3401 | 462.3479 | Positive | RP | Lipids | Sphingolipids | Sphingoid bases | Sphingoid base analogs |

| | | | | | | | | | |
|---|---|---|---|---|---|---|---|---|---|
| C13915 | Hexadecasphinganine | 273.2663 | 274.2735 | Positive | RP | Lipids | Sphingolipids | Sphingoid bases | Sphingoid base homologs and variants |
| C01530 | Octadecanoic acid | 284.2724 | 285.2739 | Positive | RP | Lipids | Fatty acyls | Fatty Acids and Conjugates | Straight chain fatty acids |
| C00163 | Propanoate | 74.0367 | 75.0446 | Positive | RP | Lipids | Fatty acyls | Fatty Acids and Conjugates | Straight chain fatty acids |
| C01585 | Hexanoic acid | 116.0833 | 117.0911 | Positive | RP | Lipids | Fatty acyls | Fatty Acids and Conjugates | Straight chain fatty acids |
| C00106 | Uracil | 112.0262 | 111.0189 | Negative | Both | Nucleic acids | Bases | Pyrimidines | NA |
| C00214 | Thymidine | 242.0903 | 241.0830 | Negative | HILIC | Nucleic acids | Nucleosides | Deoxyribonucleosides | NA |
| C00299 | Uridine | 244.0695 | 243.0622 | Negative | Both | Nucleic acids | Nucleosides | Ribonucleosides | NA |
| C00212 | Adenosine | 267.0958 | 266.0885 | Negative | HILIC | Nucleic acids | Nucleosides | Ribonucleosides | NA |
| C00294 | Inosine | 268.0811 | 267.0738 | Negative | Both | Nucleic acids | Nucleosides | NA | NA |
| C00526 | Deoxyuridine | 228.0747 | 227.0672 | Negative | HILIC | Nucleic acids | Nucleosides | Deoxyribonucleosides | NA |
| C00242 | Guanine | 151.0485 | 150.0411 | Negative | Both | Nucleic acids | Bases | Purines | NA |
| C00475 | Cytidine | 243.0855 | 242.0782 | Negative | Both | Nucleic acids | Nucleosides | Ribonucleosides | NA |
| C00380 | Cytosine | 111.0427 | 110.0348 | Negative | Both | Nucleic acids | Bases | Pyrimidines | NA |
| C00178 | Thymine | 126.0424 | 125.0346 | Negative | HILIC | Nucleic acids | Bases | Pyrimidines | NA |
| C00881 | Deoxycytidine | 227.0910 | 226.0832 | Negative | HILIC | Nucleic acids | Nucleosides | Deoxyribonucleosides | NA |
| C00387 | Guanosine | 283.0926 | 282.0847 | Negative | HILIC | Nucleic acids | Nucleosides | Ribonucleosides | NA |
| C00147 | Adenine | 135.0544 | 136.0616 | Positive | RP | Nucleic acids | Bases | Purines | NA |
| C00122 | Fumarate | 116.0098 | 115.0026 | Negative | HILIC | Organic acids | Carboxylic acids | Dicarboxylic acids | NA |
| C00042 | Succinate | 118.0255 | 117.0182 | Negative | HILIC | Organic acids | Carboxylic acids | Dicarboxylic acids | NA |
| C00022 | Pyruvate | 88.0149 | 87.0076 | Negative | HILIC | Organic acids | Carboxylic acids | 2-Oxocarboxylic acids | NA |
| C00311 | Isocitrate | 192.0269 | 191.0191 | Negative | HILIC | Organic acids | Carboxylic acids | Tricarboxylic acids | NA |
| C00763 | D-Proline | 115.0635 | 116.0707 | Positive | HILIC | Amino acids | Peptides | Other amino acids | NA |
| C00183 | L-Valine | 117.0790 | 118.0863 | Positive | HILIC | Amino acids | Peptides | Common amino acids | NA |
| C00079 | L-Phenylalanine | 165.0785 | 166.0858 | Positive | Both | Amino acids | Peptides | Common amino acids | NA |
| C00041 | L-Alanine | 89.0476 | 90.0554 | Positive | Both | Amino acids | Peptides | Common amino acids | NA |
| C00483 | Tyramine | 137.0833 | 138.0911 | Positive | HILIC | Amines | Peptides | Biogenic amines | NA |
| C00082 | L-Tyrosine | 181.0734 | 182.0812 | Positive | HILIC | Amino acids | Peptides | Common amino acids | NA |
| C01879 | 5-Oxoproline | 129.0415 | 128.0342 | Negative | Both | Amino acids | Peptides | Other amino acids | NA |
| C00407 | L-Isoleucine | 131.0935 | 130.0863 | Negative | HILIC | Amino acids | Peptides | Common amino acids | NA |
| C00148 | L-Proline | 115.0623 | 182.0428 | Negative | Both | Amino acids | Peptides | Common amino acids | NA |
| C00025 | L-Glutamate | 147.0522 | 146.0449 | Negative | Both | Amino acids | Peptides | Common amino acids | NA |

| | | | | | | | | | |
|---|---|---|---|---|---|---|---|---|---|
| C00188 | L-Threonine | 119.0572 | 186.0377 | Negative | Both | Amino acids | Peptides | Common amino acids | NA |
| C00327 | L-Citrulline | 175.0951 | 174.0877 | Negative | HILIC | Amino acids | Peptides | Other amino acids | NA |
| C00065 | L-Serine | 105.0420 | 104.0342 | Negative | HILIC | Amino acids | Peptides | Common amino acids | NA |
| C00791 | Creatinine | 113.0584 | 112.0506 | Negative | HILIC | NA | Peptides | NA | NA |
| C00152 | L-Asparagine | 132.0530 | 131.0452 | Negative | HILIC | Amino acids | Peptides | Common amino acids | NA |
| C00077 | L-Ornithine | 132.0893 | 131.0815 | Negative | HILIC | Amino acids | Peptides | Other amino acids | NA |
| C00049 | L-Aspartate | 133.0370 | 132.0292 | Negative | HILIC | Amino acids | Peptides | Common amino acids | NA |
| C00064 | L-Glutamine | 146.0687 | 145.0609 | Negative | Both | Amino acids | Peptides | Common amino acids | NA |
| C00073 | L-Methionine | 149.0506 | 148.0428 | Negative | Both | Amino acids | Peptides | Common amino acids | NA |
| C00606 | 3-Sulfino-L-alanine | 153.0092 | 152.0013 | Negative | HILIC | Amino acids | Peptides | Other amino acids | NA |
| C00135 | L-Histidine | 155.0704 | 154.0626 | Negative | HILIC | Amino acids | Peptides | Common amino acids | NA |
| C00078 | L-Tryptophan | 204.0899 | 203.0821 | Negative | Both | Amino acids | Peptides | Common amino acids | NA |
| C16138 | L-Pyrrolysine | 255.1618 | 256.1691 | Positive | RP | Amino acids | Peptides | Other amino acids | NA |
| C00037 | Glycine | 75.0320 | 76.0398 | Positive | RP | Amino acids | Peptides | Common amino acids | NA |
| C03194 | (R)-1-Aminopropan-2-ol | 75.0684 | 76.0762 | Positive | RP | Amines | Peptides | Biogenic amines | NA |
| C00134 | Putrescine | 88.0999 | 89.1077 | Positive | RP | Amines | Peptides | Biogenic amines | NA |
| C03406 | N-(L-Arginino)succinate | 290.1213 | 291.1291 | Positive | RP | Amino acids | Peptides | Other amino acids | NA |
| C00385 | Xanthine | 152.0313 | 153.0391 | Positive | HILIC | Phytochemical compounds | Alkaloids | Others | Purine alkaloids |
| C00262 | Hypoxanthine | 136.0375 | 135.0302 | Negative | HILIC | Phytochemical compounds | Alkaloids | Others | Purine alkaloids |
| C00253 | Nicotinate | 123.0315 | 124.0393 | Positive | Both | Vitamins and Cofactors | Vitamins | Water-soluble vitamins | NA |
| C00061 | FMN | 456.1097 | 455.1025 | Negative | HILIC | Vitamins and Cofactors | Cofactors | Coenzymes | NA |
| C00847 | 4-Pyridoxate | 183.0531 | 182.0452 | Negative | HILIC | Vitamins and Cofactors | Vitamins | Water-soluble vitamins | NA |
| C00864 | Pantothenate | 219.1109 | 218.1031 | Negative | Both | Vitamins and Cofactors | Vitamins | Water-soluble vitamins | NA |
| C00053 | 3'-Phosphoadenylyl sulfate | 506.9860 | 505.9782 | Negative | HILIC | Vitamins and Cofactors | Cofactors | Coenzymes | NA |
| C00018 | Pyridoxal phosphate | 247.0247 | 248.0259 | Positive | RP | Vitamins and Cofactors | Cofactors | Coenzymes | NA |
| C00153 | Nicotinamide | 122.0475 | 123.0553 | Positive | RP | Vitamins and Cofactors | Vitamins | Water-soluble vitamins | NA |
| C00250 | Pyridoxal | 167.0575 | 168.0653 | Positive | RP | Vitamins and Cofactors | Vitamins | Water-soluble vitamins | NA |
| C00072 | Ascorbate | 176.0346 | 177.0425 | Positive | RP | Vitamins and Cofactors | Vitamins | Water-soluble vitamins | NA |

| | | | | | | | | | |
|---|---|---|---|---|---|---|---|---|---|
| C09714 | Riboflavin | 376.1833 | 377.1457 | Positive | RP | Vitamins and cofactors | Vitamins | Water-soluble vitamins | NA |
